# Neonatal BCG Vaccination Engages the Vasculature to Elicit γδ T Cell–Mediated Protec-tion against Tuberculosis

**DOI:** 10.64898/2025.12.08.693020

**Authors:** Mina Sadeghi, Nargis Khan, Yao Chen, Leonardo F. Jurado, Anshul Sinha, Katalina Couto, Ian Vong, Veronica Locher, Kim A. Tran, Shahin Bolori Hanafi, Bavanitha Thuraira-jah, Tanishia Jean Shiri, Nisreen Ghanem, Eva Kaufmann, Jia Ming Low, Zubair Amin, Corinne Maurice, Julia Kolter, Jianguo Xia, Mikael Knip, Ramnik Xavier, Mihai G. Netea, Irah King, Luis B. Barreiro, Ajitha Thanabalasuriar, Amit Singhal, Maziar Divangahi

## Abstract

The Bacillus Calmette–Guérin (BCG) vaccine remains the only approved vaccine against tuberculosis (TB). Although its efficacy against pulmonary TB in adults is limited, BCG provides remarkable protection against miliary TB when administered during infancy. Despite more than 100 million infants worldwide receiving BCG annually, the mechanisms underlying its neonatal protective effects remain poorly defined. Here, we demonstrate that subcutaneous neonatal BCG vaccination (BCG-sc) induces a marked expansion of γδ T cells producing IL-17 and IL-22, which mediated protection against subsequent *Mycobacterium tuberculosis* (*Mtb*) experimental infection. A similar expansion of γδ T cells was observed in a longitudinal cohort of infants, from birth to three months of infants followed after intradermal BCG vaccination. Mechanistically, BCG-mediated protection in neonates was linked to its early vascular dissemination through the distinct structure of neonatal skin, resembling the protective effects of intravenous BCG in adults. Moreover, neonatal BCG-sc vaccination generated a distinct BCG-induced microbiome signature, characterized by enrichment of *Prevotellaceae*, *Tannerellaceae*, and *Bifidobacteriaceae*, which was associated with protection. Together, these findings identify γδ T cells as key mediators of early-life BCG-induced immunity and highlight the role of the gut–lung axis in long-term protection against TB from infancy into adulthood.

## Introduction

*Mycobacterium tuberculosis* (*Mtb*) has persisted as the leading cause of death from infectious disease worldwide, claiming approximately 1.5 million lives annually^1^. Despite more than a century of research, the Bacillus Calmette–Guérin (BCG) vaccine remains the sole licensed vaccine available against tuberculosis (TB)^2^. Neonatal intradermal BCG vaccination confers approximately 70% protection against miliary and meningeal TB in children, but its efficacy in adults ranges widely from 0–80%^3^. Strikingly, in adult non-human primates (NHPs), systemic (intravenous; IV) BCG administration provides markedly enhanced protection, and in some cases sterilizing immunity, compared to the intradermal (ID) route^4^. This superior protection has been associated with increased systemic T cell responses^4, 5^. In an adult mouse model, intravenous BCG (BCG-iv) has also been shown to rapidly access the bone marrow, reprogramming hematopoietic stem cells (HSCs) to generate innate memory responses (trained immunity) against TB^6^, a program that *Mtb* actively suppresses^7^. Given that more than 100 million infants worldwide receive BCG-ID within the first week of life^2^, and that neonatal BCG provides robust protection, elucidating its protective mechanisms in infants will be critical for defining the cellular and molecular basis needed to develop next-generation TB vaccines.

There are three fundamental differences between neonates and adults that may underlie the enhanced protective effects of BCG-ID vaccination in early life. First, neonatal skin differs structurally and compositionally from adult skin, with a thinner stratum corneum and epidermis^8^, which may facilitate early dissemination of BCG. Second, the neonatal immune system is distinct not merely due to immune inexperience^9^, but also because of its unique cellular composition. At mucosal sites, neonates harbor a higher relative abundance of innate-like γδ T cells, whereas αβ T cells are comparatively sparse^10, 11^. Third, the neonatal gut microbiome is highly dynamic in early life, before stabilizing in adulthood^12^. For example, an increased abundance of *Bifidobacterium* during infancy correlates with stronger immune responses to BCG vaccination and long-term effects on immunity^13^, underscoring the direct contribution of the gut microbiome to the durability of BCG-induced protection.

In this study, we tested these three possibilities to investigate the protective mechanism of BCG vaccination in neonates. We found that BCG-sc vaccination in neonates mimics the effects of BCG-iv in adults by accessing the neonatal skin vasculature and enabling rapid systemic dissemination to other organs. In both mice and humans, this resulted in a robust expansion of γδ T cells, which were essential for protection against TB. In the murine model, these γδ T cells produced high amounts of IL-17A and IL-22, conferring long-lasting protection against *M. tuberculosis* into adulthood, a process associated with a distinct BCG-induced microbiome signature.

## Results

### Neonatal BCG vaccination offers long-lasting protection

To compare the effects of BCG vaccination in neonates and adults, we vaccinated neonatal C57BL/6 mice at post-natal days (PND) 2-3 or adult mice between 6-8 weeks old via subcutaneous (BCG-sc) or intravenous (BCG-iv) routes at the same dose of 1 x10^6^ CFU of BCG (Tice strain) (**Fig. 1a**). After six weeks of BCG-vaccination, the mice were infected with *Mtb* (H37Rv; ∼50 CFU) and following two weeks *Mtb* burdens and histopathology of the lungs were assessed (**Fig. 1b**). Neonatal mice exhibited a significant reduction in pulmonary bacterial burdens after BCG-sc or BCG-iv compared to PBS control, while adult mice demonstrated significant protection after BCG-iv (**Fig. 1b**). Hence, the protection against Mtb between BCG-sc and BCG-iv in neonates was almost identical. Following four weeks of *Mtb* infection, lung histology demonstrated fewer pathological granuloma-like nodes and a lower histopathological score in the lungs of neonatal mice compared to the PBS control and adult counterparts (**Fig. 1c-d**). To test the duration of BCG-mediated protection after vaccination in neonatal mice compared to adults, mice were rested 20 weeks (**Fig. 1e-g**) post-BCG vaccination. Four weeks after aerosolized infection with *Mtb*, neonatally vaccinated mice displayed significant reductions in bacterial burden in both the lung (**Fig. 1f**) and spleen (**Fig. 1g**), while BCG-protection was lost in mice vaccinated at adult age. Given these findings, neonatal mice alone were further aged one-year post-BCG vaccination and infected with *Mtb*-aerosol (**Fig. 1h**). BCG-vaccination in neonatal mice was still able to reduce bacterial burdens in the lungs and spleen (**Fig. 1i-j**). Thus, BCG vaccination in murine neonates recapitulates the epidemiological observations from human neonatal BCG vac-cination.

**Figure 1:**
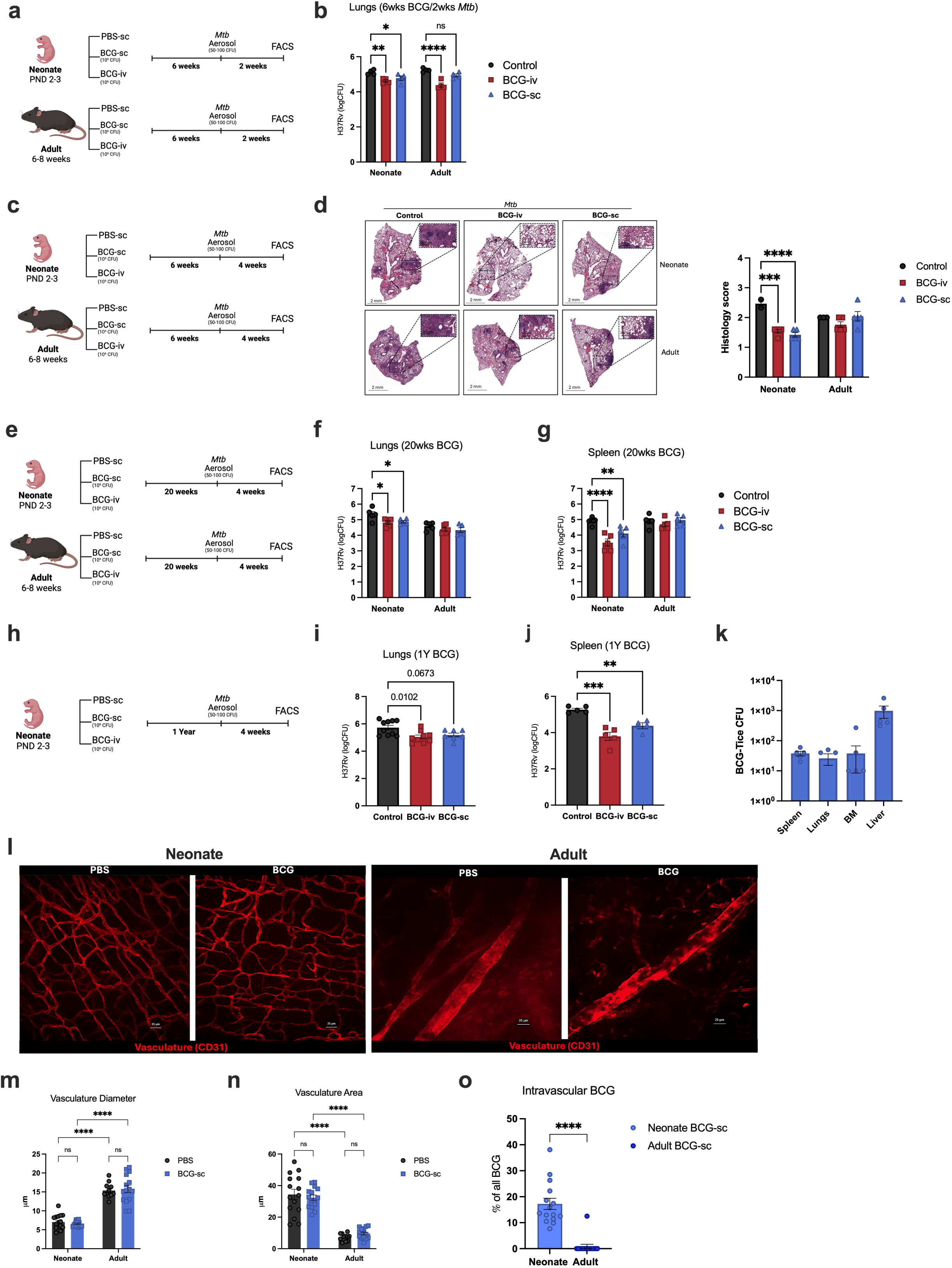
Neonatal BCG vaccination provides strong protection against *Mtb* infection and leads to systemic dissemination. **(a)** Neonates at PND 2-3 or adult C57BL/6 mice at 6-8 weeks were vaccinated by subcutaneous or intravenous routes or administered PBS. 6 weeks post-vaccination, mice were challenged with aerosolized Mtb-infection (H37Rv, 50-100 CFU). Two weeks following infection, H37Rv burden was assessed. **(b)** H37Rv lung CFU in neonates or adults two weeks following *Mtb*-aerosol challenge. N=5-6 per group. **(c)** Experimental model and **(d)** representative H&E (1X) images (left) for granulomatous lesions in lung lobes of neonatal (top) or adult (bottom) mice one month following *Mtb*-aerosol infection with quantification (right, N=5 per group). **(e)** Experimental model for vaccination of neonates and adults. Mice were infected with *Mtb*-aerosol infection 20 weeks following initial vaccination. H37Rv CFU of **(f)** lungs or **(g)** spleen in neonates or adults, 20 weeks post-BCG vaccination and one month following *Mtb*-aerosol challenge. N=5-6 per group. **(h)** Experimental model for vaccination of neonates and adults. Mice were infected with *Mtb*-aerosol infection 1 year following BCG vaccination. H37Rv CFU of **(i)** lungs or **(j)** spleen in neonates or adults, 1 year post-BCG vaccination and one month following *Mtb*-aerosol challenge. N=7-10 per group. **(k)** BCG-Tice CFU in spleens, whole lungs, femur bone marrow and liver of neonatal mice 6 hours post BCG vaccination. N=5 per group. **(l)** Wholemount immunofluorescence staining of CD31+ vasculature in neonate (left) and adult (right) skin with PBS-sc or BCG-sc injections, 5 minutes post-injection. (Scale bar: 25 μm) **(m)** Average vessel diameter was calculated by measuring 40 individual vascular segments across the entire field of view (FOV). **(n)** Vascular area was quantified by calculating the proportion of CD31-positive area relative to the total area of the FOV. **(o)** Three-dimensional reconstruction of wholemount imuunofluorescence from BCG-vaccinated neonate and adult mice, rendered using Imaris software. Scale bar: 10 μm. **(p)** Intravascular BCG was calculated by using the Imaris software to quantify the frequency of BCG in the vasculature over total BCG per FOV. For all wholemount immunoflorescence, n=3 PBS treated neonates, n=3 BCG vaccinated (7x 10^6^ CFU) neonates, n=3 PBS treated adult, n=3 BCG vaccinated (7x 10^6^ CFU) adults. Five randomized technical replicates (FOV) were taken per biological replicate. Statistical significance for comparison between neonates and adults were performed by a Mann-Whitney t-test (**m-o**). One (**i, j, k**) or two-way (**b, d, f, g)** ANOVA followed by Tukey’s multiple comparisons test, data are presented as Mean ± SEM with * p < 0.05, ** p < 0.01, *** p < 0.001, **** p < 0.0001.

### BCG disseminates through the vascular bed of neonatal but not adult skin

Considering the identical protection between BCG-iv and BCG-sc vaccination in neonates (**Fig. 1b, f, i**), we postulated that BCG-sc vaccination may lead to systemic dissemination of BCG in neonates. Interestingly, as early as 24 hours post neonatal BCG-sc vaccination, BCG colonies were disseminated and measured in the spleen, lungs, bone marrow (BM) and liver (**Fig. 1k**). Given that the skin is the first portal for the vaccine, and the well-reported differences in the structure of skin tissue through early development^8^, we leveraged whole-mount immunofluorescence microscopy to juxtapose the structure and vascularization of neonatal and adult skin immediately following BCG vaccination. In neonates and adults, there were no significant differences in the vasculature structure, as measured by CD31 staining, between PBS controls and BCG-sc groups (**Fig. 1l**). While adults displayed higher vessel diameters than neonates (**Fig. 1l-m**), the vasculature area, which is the frequency of vessels over the total area of the field of view (FOV), was significantly increased in neonates compared to adults (**Fig. 1l, n**). The observation of increased vasculature area in neonates compared to adults was further replicable in another independent laboratory (**Extended Data Fig. 1a-b**). These data demonstrate that within neonatal skin, the vasculature forms a more intricate network of smaller vessels, while the adult skin is composed of fewer, larger vessels. In BCG-sc neonates by PND7, CD31 staining demonstrated an intermediate phenotype; both small intricate vessels as well as large vessels were observed within the same FOV (**Extended Data Fig. 1c**) with increased vasculature diameters, but similar vasculature area compared to PND3 neonates (**Extended Data Fig. 1d-e**). Next, we sought to quantify the intravascular abundance of BCG between neonates and adults. Three-dimensional reconstruction of skin samples demonstrated an increased frequency of intravascular BCG over total BCG per FOV within neonatal skin **(Supplementary Video 1)** compared to adult skin (**Supplementary Video 2, Fig 1o-p**). By Z-plane optical sectioning of adult murine skin, it was observed that the majority of BCG were located within the adipose tissue layer compared to regions adjacent to the vascular network (**Supplementary Video 3)**. In fact, for the quantification of vascular diameter and area in adult mice, the process of adipose tissue removal eliminated most detectable BCG. Together, these data demonstrate that BCG readily enters the neonatal vasculature, whereas in adults, the thick layer of subcutaneous adipose tissue leads to entrapment of BCG colonies and prevents dissemination into the vasculature. Thus, in the neonatal mouse, the subcutaneous vaccination strategy mirrors the dissemination kinetics of the intravenous administration.

### Neonatal BCG vaccination induces **γδ** T cell expansion

One key feature of intravenous BCG vaccination in adults is the direct access to BM hematopoietic stem cells (BM-HSCs) and the induction of central trained immunity (Kaufmann et al. Cell 2018;). To interrogate the induction of central training, we assessed BM HSCs after four weeks of neonatal BCG-sc vaccination (**Fig. 2a and Supplementary Figure 1**). Neonatal vaccination induced a significant increase in LKS^+^ HSCs (**Fig. 2b**), including both short term (ST) HSCs and multipotent progenitors (MPPs; **Extended Data Fig. 2a-h**). In contrast to BCG-iv vaccination in adults^6^, the common myeloid progenitor (CMP) was significantly reduced post-vaccination, while the common lymphoid progenitor (CLP) displayed an expansion (**Fig. 2c-d**). Thus BCG-sc vaccination in neonatal hematopoiesis induced an expansion of HSPCs with the bias towards lymphopoiesis.

**Figure 2:**
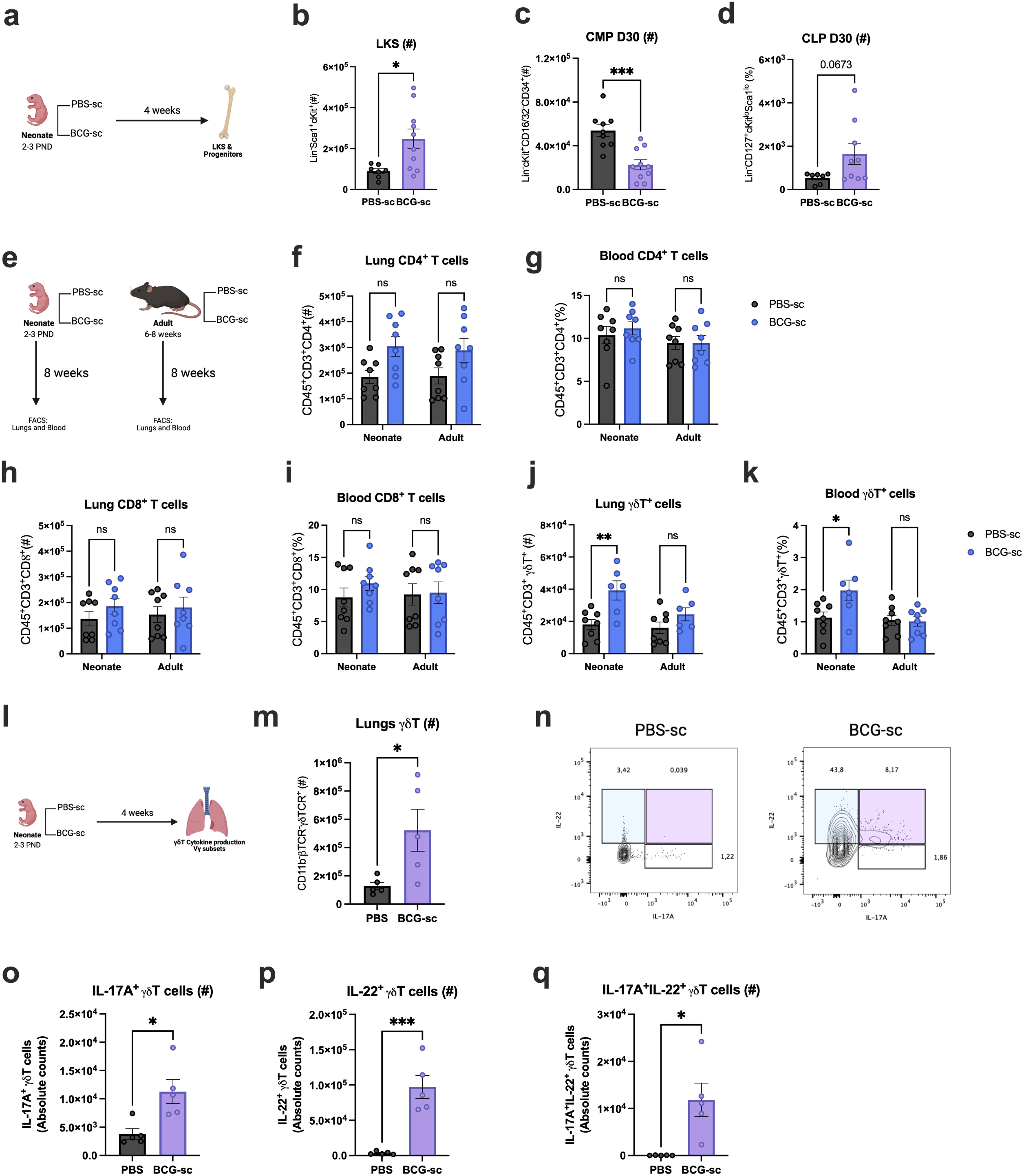
Neonatal BCG vaccination led to increased γδ T cells in the lungs and blood. **(a)** Experimental model for BCG-sc vaccination of neonates and collection of BM following 4 weeks. Absolute counts of **(b)** LKS^+^ cells, **(c)** CMPs, and **(d)** CLPs in BM of neonatal mice 28 days post-administration of PBS-sc or BCG-sc (N= 7-10 per group). **(e)** Mice were vaccinated with BCG-sc at PND 2-3 (neonates) or 6-8 weeks (adults) and lung and blood samples were collected after 8 weeks. **(f)** Absolute counts of lung CD4^+^ T cells and **(g)** frequencies of blood CD4^+^ T cells. **(h)** Absolute counts of CD8^+^ T cells and **(i)** frequencies of blood CD8^+^ T cells. **(j)** Abso-lute counts of Y ^+^ T cells and **(k)** frequencies of blood γδ T cells. (N= 8 per group). **(l)** Experimental design for BCG-sc vaccination of neonates and collection of lungs following 4 weeks. **(m)** Absolute counts and **(n)** FACS plots of pulmonary CD11b^-^ CD3^+^ βTCR^-^ γδ TCR^+^ T cells **(o)** IL-17A^+^ γδ T cells, **(p)** IL-22^+^ γδT cells, and **(q)** IL- 17A^+^IL-22^+^ γδ T cells at 28 days post- vaccination (N=5 per group). Data are presented as mean ±SEM and are representative of two independent experiments. Statistical significance for comparison were performed by Two-way ANOVA (**f-k**) followed by Sidak’s multiple comparisons test or unpaired two-tailed t-test (**b, c, d, m, o, p, q**), data are presented as Mean ± SEM with * p < 0.05, ** p < 0.01.

We next immunophenotyped mature innate and adaptive immune cells in the blood and lung of neonates and adults at eight weeks post-BCG-sc vaccination (**Fig. 2f and Supplementary Figure 1**). While there were no significant differences in most innate and adaptive cell types (**Extended Data Fig. 2i-p**), including conventional CD4^+^ and CD8^+^ T cells (**Fig. 2f-i**), there was a significant increase in the frequency and absolute counts of γδ T cells in the blood and lungs of neonatal mice vaccinated with BCG-sc (**Fig. 2j-k**). Unconventional T cells predominate during early life; in mice, γδTCR expression is first detectable at gestational day 14 and in humans γδ T cells develop within the fetal thymus and consist primarily of Vδ2 T cells, which predominate after birth^10, 14^. We next further characterized unstimulated γδ T cells at four weeks post-BCG vaccination in the lung (**Fig. 2l**) and demonstrated a five-fold increase in the number of γδ T cells in neonatal lungs (**Fig. 2m**). This expansion was associated with an increased production of interleukin (IL)-17A and IL-22 by γδ T cells (**Fig. 2n-p**), as well as double positive IL-17A^+^IL-22^+^ γδ T cells (**Fig. 2q**), but not IFNγ, IL-4, or IL-13 (**Extended Figure 3a-c**). Together, neonatal BCG-sc vaccination promoted a shift toward lymphopoiesis in the BM and expanded pulmonary γδ T cells, which produced increased amounts of IL-17A and IL-22, both cytokines essential for protective immunity against TB^15, 16, 17^.

### BCG-induced **γδ** T cells mediate protection against *Mtb* infection

Early expansions in γδ T cells have been well documented following BCG vaccination or *Mtb* infection in both NHP models and human studies^15, 18, 19, 20, 21^. To further define their early dynamics, we employed a six-week neonatal BCG-sc vaccination model followed by *Mtb* aerosol infection (**Fig. 3a**). Four weeks after *Mtb* challenge, neonatally BCG-sc–vaccinated mice showed no differences in αβ T cells (**Fig. 3b**), including CD4 and CD8 subsets, nor in other innate or adaptive cell populations (**Extended Data Fig. 4a–h**). However, they displayed a significant increase in γδ T cells (**Fig. 3c**), with increased proliferative capacity as measured by Ki67 incorporation (**Fig. 3d**). Consistent with our findings from BCG-sc vaccination alone, γδ T cells from BCG-sc–vaccinated and *Mtb*-infected neonates exhibited elevated IL-17A (**Fig. 3e**) and IL-22 (**Fig. 3f**) production. The predominant subset was Vγ6 γδ T cells (**Fig. 4g**), rather than Vγ4, Vγ3/5, or Vγ1 subsets (**Extended Data Fig. 4i–k**). To extend these observations, we evaluated mice 20 weeks after neonatal or adult BCG-sc vaccination followed by *Mtb* aerosol infection (**Fig. 3h**). One week post-infection, BCG-sc–vaccinated neonates displayed a significant expansion of pulmonary γδ T cells compared to PBS controls or adult BCG-sc–vaccinated mice (**Fig. 3i**). By five weeks post-infection, neonatally vaccinated mice continued to maintain significantly higher levels of total γδ T cells than their adult-vaccinated counterparts (**Fig. 3j**). Collectively, these findings demonstrate that neonatal BCG-sc vaccination promotes protection against Mtb infection possibly through the expansion of Vγ6 IL-17A IL-22 γδ T cells.

**Figure 3:**
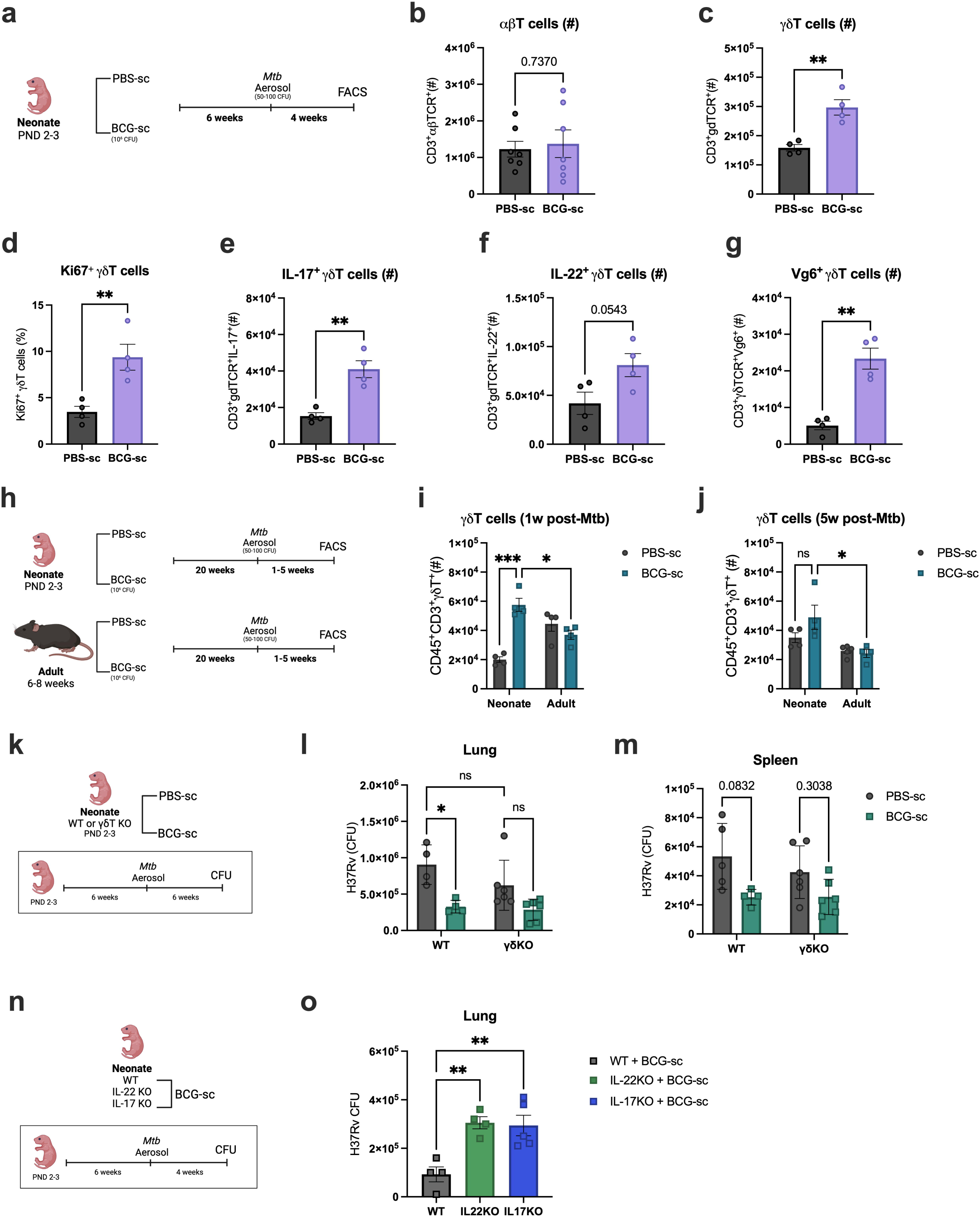
Neonatal BCG-sc induced γδT cells were necessary for protection against *Mtb*-infection. **(a)** Experimental model of neonatal BCG-sc vaccination for 6 weeks, followed by *Mtb*-aerosol infection and analysis after four weeks. Absolute counts of pulmonary **(b)** αβ T cells (N= 8 per group), **(c)** γδ T cells, **(d)** Ki67^+^ γδ T cells, **(e)** IL-17A^+^ γδ T cells, **(f)** IL-22^+^ γδT cells, and **(g)** Vγ6^+^ γδ T cells (N=4 per group). **(h)** Experimental model of neonatal and adult BCG vaccination for 20 weeks, followed by *Mtb*-aerosol challenge (50-100 CFU). Absolute counts of pulmonary γδ T cells following **(i)** one week or **(j)** five weeks of *Mtb*-infection (N=4-5 per group). **(k)** Experimental model for neonatal vaccination in WT C57BL/6 or δTCR^-/-^C57BL/6 mice. After 6 weeks of neonatal BCG-sc vaccination, mice were infected with Mtb-aerosol, and assessed after 6 weeks post-infection. H37Rv CFU in the **(l)** lungs and **(m)** spleen of WT or γδTCR^-/-^ neonatal mice (N=4-6 per group). **(n)** Experimental model for neonatal vaccina-tion in WT C57BL/6, IL-22^-/-^ C57BL/6, or IL-17^-/-^ C57BL/6 mice. **(o)** H37Rv CFU in the lungs and of WT, IL-22^-/-^ or IL-17^-/-^ neonatal mice. N=4-5 per group. Data are representative of two independent experiments. Statistical significance for comparison were performed by one (**o**) or two-way (**i, j, l, m**) ANOVA followed by Sidak’s (**i, j)** or Tukey’s (**l, m, o**) multiple comparisons test or unpaired two-tailed t-test (**b-g**), data are presented as Mean ± SEM with * p < 0.05, ** p < 0.01, *** p < 0.001.

**Figure 4:**
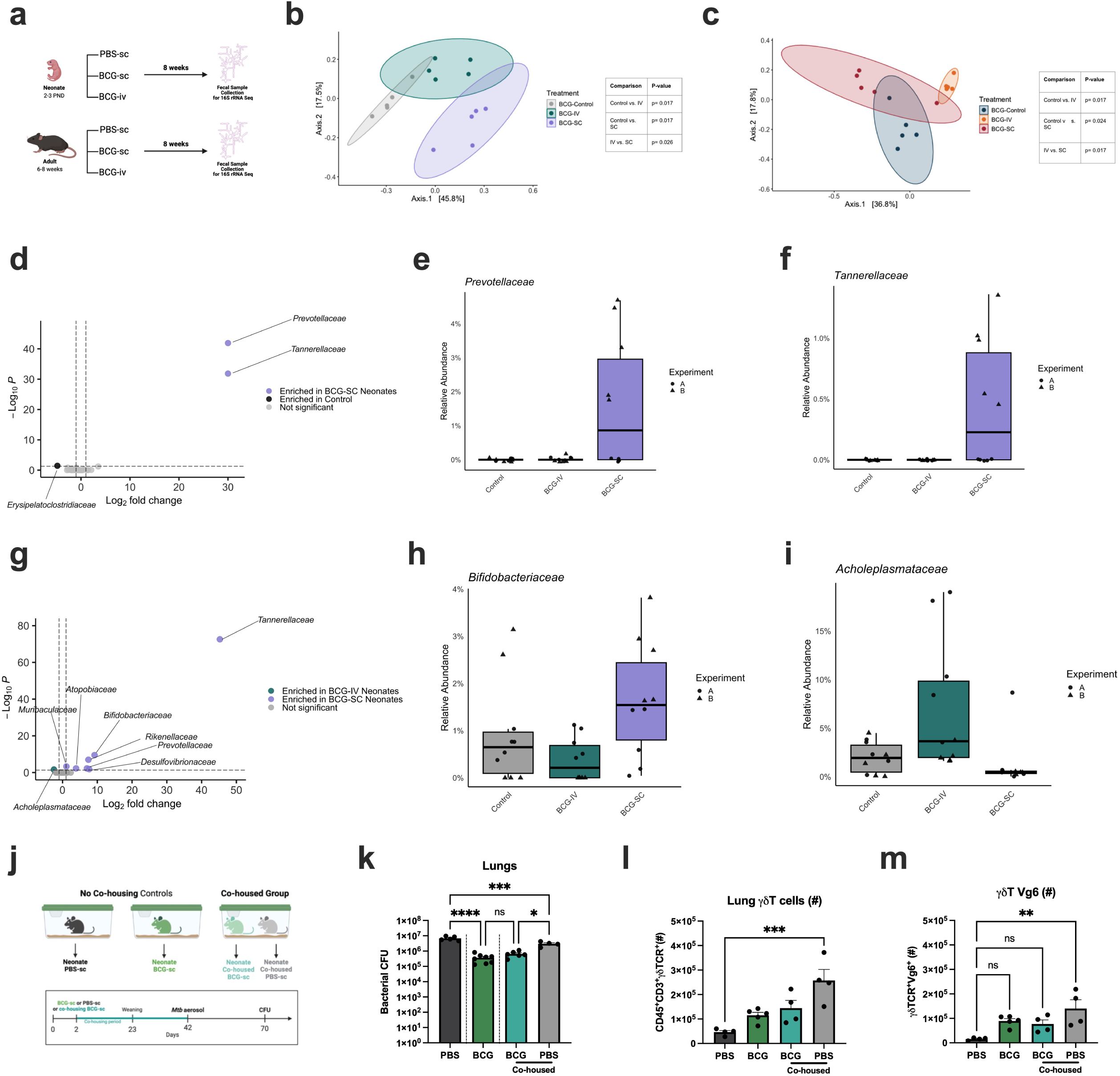
BCG-sc vaccinated neonates display a unique and protective early-life micro-biome signature. **(a)** Experimental model for DNA extraction and 16S rRNA sequencing of fe-cal samples from adult and neonate mice. Principal (N=5 per group). Principal Coordinates Analysis (PCoA) with Bray-Curtis dissimilarity for **(b)** neonatal mice or **(c)** adult mice. PER-MANOVA was used to assess significance p < 0.05 between groups. These data are representa-tive of one of two independent experiments. **(d)** DESeq2 analysis for enriched bacterial taxa be-tween neonatal BCG-sc and PBS controls. Relative abundance of neonatal **(e)** *Prevotellaceae* and **(f)** *Tannerellaceae* families in neonatal groups. **(g)** DESeq2 analysis for enriched bacterial taxa between BCG-sc and BCG-iv treatments. Log_2_ fold-change > 1 or < -1 and p ≤ 0.05 were used as thresholds for significance. Relative abundance of **(h)** *Bifidobacteriaceae* and **(i)** *Achole-plasmataceae* families in neonatal groups. **(j)** Experimental set-up for co-housing study of neo-natal mice vaccinated with BCG-sc or PBS-sc controls. Mice in the co-housed group were co-housed for approximately 40 days from vaccination date at 2 PND. After the co-housing or expo-sure period, mice were challenged with *Mtb*-aerosol infection (H37Rv, 50-100 CFU). Mice were assessed one month following aerosolization. Data are representative of three independent ex-periments. **(k)** Lung H37Rv CFU of no-cohousing controls (dark grey, green) or cohoused mice vaccinated with BCG-sc (teal) or PBS control (light grey) (N=4-5 per group). Absolute counts of pulmonary **(l)** γδ T cells and **(m)** Vγ6^+^ γδ T cells. N=4-5 per group. One way ANOVA (**k-m**) fol-lowed by Tukey’s multiple comparisons test, data are presented as Mean ± SEM with * p < 0.05, ** p < 0.01, *** p < 0.001, **** p < 0.0001.

To determine the direct role of γδ T cells in BCG-sc–mediated protection against Mtb in neonates, wild-type (WT) and TCRδ / (γδ T cell–deficient) neonatal mice were vaccinated with BCG-sc and challenged with aerosolized *Mtb* six weeks later (**Fig. 3k**). Bacterial burdens assessed six weeks post-infection revealed that BCG-sc–vaccinated TCRδ / neonates failed to show protection in the lungs (**Fig. 3l**) or spleen (**Fig. 3m**) compared to unvaccinated PBS-sc controls. Since IL-22^16, 22^ and IL-17^17, 23^ have been reported to play a protective role against TB, and γδ T cells from BCG-sc–vaccinated neonates produced significant amounts of IL-22 and IL-17, we next investigated its contribution to protection. WT, IL-22 /, and IL-17 / neonatal mice were vaccinated with BCG-sc, and four weeks later exposed to aerosolized *Mtb* (**Fig. 3n**). Four weeks post-infection, BCG-sc–vaccinated IL-22 / and IL-17 / mice showed a loss in protection compared to BCG-sc-vaccinated WT mice (**Fig. 3o**). In summary, neonatal BCG-sc vaccination induces γδ T cells that are required for protection against *Mtb*, with IL-22 and IL-17 playing a critical role for this protection.

### A protective neonatal BCG microbiome signature

A growing body of literature has demonstrated that during early life the gut microbiome imparts important influence on the abundance and function of immune cells at barrier tissues^24^, particularly unconventional T cells^25^. Based on this, in two independent experiments, we assessed the composition of the fecal microbiome of neonatally- or adult-vaccinated animals with BCG-sc, BCG-iv or PBS controls by 16S rRNA gene sequencing (**Fig. 4a**). We observed that the experiment (PERMANOVA p= 0.001, R^2^=0.15) and the age of the mouse (PERMANOVA p= 0.001, R^2^=0.10711) had significant effects on bacterial diversity (**Extended Data Fig. 5a-b**). Given the differences in microbiome composition between experiments, we assessed the effects of BCG-sc and BCG-iv in each experiment in adults and neonates. We found that in both experiments there were treatment-specific differences in beta-diversity as measured by Bray-Curtis distance. Both BCG-sc and BCG-iv administration induced robust differences in beta diversity compared to all other groups, both within neonates (**Fig. 4b**) and adults (**Fig. 4c**), as measured by Bray-Curtis distance, which was also replicable in a second experiment (**Extended Data Fig. 5c-d**). To assess the changes in specific bacterial taxa within BCG-sc vaccinated neonates compared to PBS controls, we performed differential abundance analyses using DESeq2 to determine the uniquely enriched families (**Fig. 4d**). In BCG-sc vaccinated neonates, *Prevotellaceae* (**Fig. 4e**), and *Tannerellaceae* (**Fig. 4f**) families were increased, while *Erysipelatoclostridiaceae* was enriched in PBS controls (**Extended Data Fig. 5e**). In the comparison of BCG-sc neonates with BCG-iv neonates (**Fig. 4g**), *Prevotellaceae* and *Tannerellaceae* were again enriched in BCG-sc neonates, as well as *Desulfovibrioceae* (**Extended Data Fig. 5f**), *Rikenellaceae* (**Extended Data Fig. 5g**), *Bifidobacteriaceae* (**Fig. 4h**), *Muribaculaceae* (**Extended Data Fig. 5h**) and *Atopobiaceae* (**Extended Data Fig. 5i**). The sole enriched family unique for BCG-iv compared to BCG-sc was *Acholeplasmataceae* (**Fig. 4i**), while *Muribaculaceae* and *Atopobiaceae* were enriched in BCG-sc (Extended Data Fig.6h-i). It should be noted that some of the trends observed in the differentially abundant taxa were experiment specific. These data indicate an altered microbiome post-BCG vaccination in both adults and neonates.

**Figure 5:**
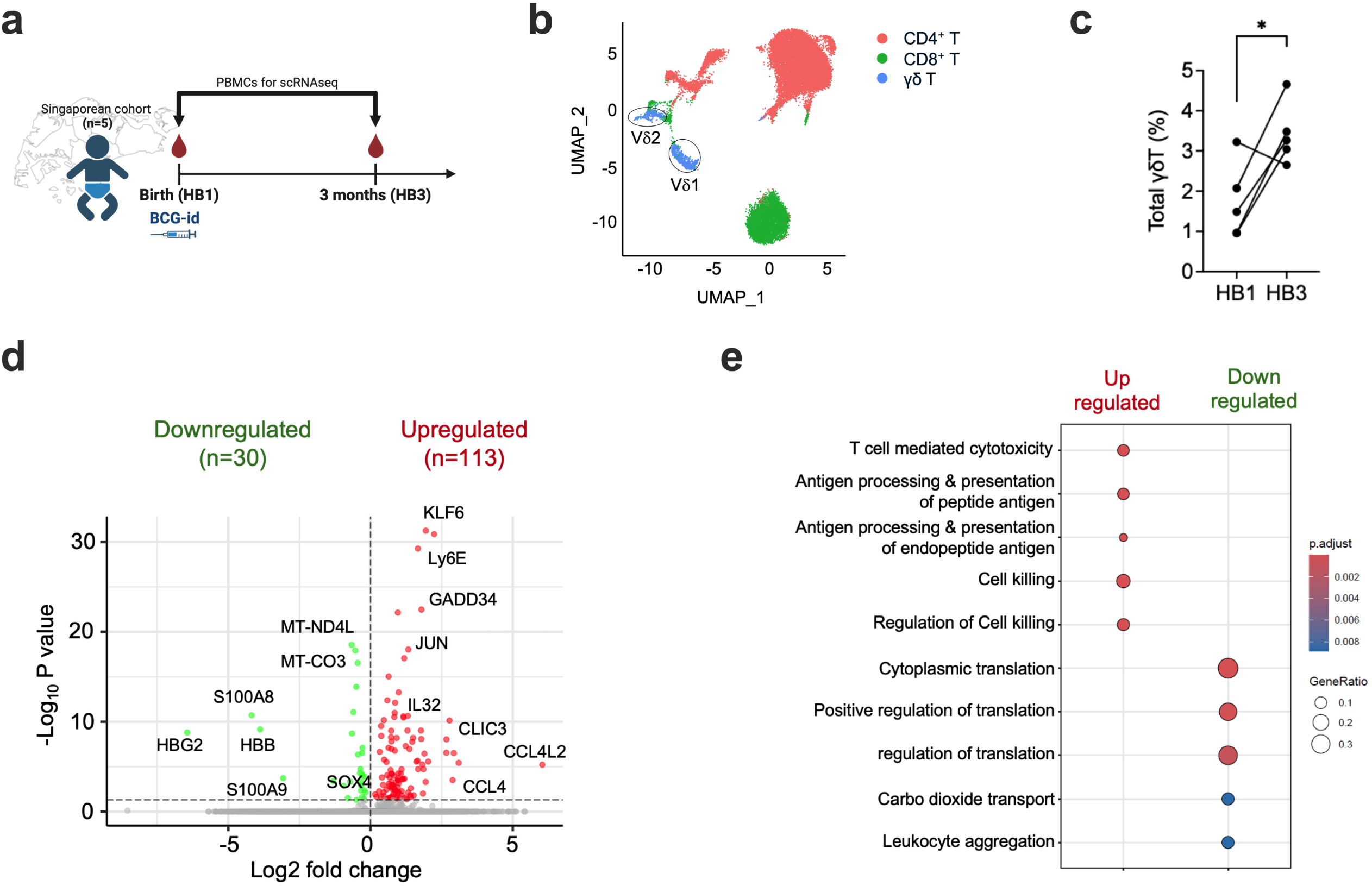
Human neonatal BCG vaccination induced expansion and transcriptional reprogramming of γδ T cells. **(a)** Study design for human neonatal blood collection at birth (HB1) and following 3 months (HB3) of BCG vaccination (N=5). **(b)** Uniform manifold approximation and projection (UMAP) plot of human neonatal T cell subtypes, including six CD4^+^ T cell sub-types, four CD8^+^ T cell subtypes, and two γδ T cell subtypes, namely Vδ1 and Vδ2 cells (circled). **(c)** Percentage of γδ T in PBMCs of infants at birth and at 3 months. * p<0.05, paired t-test. **(c)** Representative uniform manifold approximation and projection (UMAP) plots of Vδ1 and Vδ2 expression by single-cell RNA-seq in neonatal PBMCs. **(d)** Volcano plot of DEGs among γδ T cells in 3 months (HB3) vs birth (HB1). **(e)** Enriched GO biological processes in up-and down-regulated DEGs, 3 months vs birth.

To directly link the altered microbiome post-BCG-sc to the abundance and function of γδ T cells, as well as vaccine-mediated protection, we performed a co-housing study (**Fig. 4e**). In this model, there were four neonatal groups for comparison: 1) non-cohoused PBS-sc controls, 2) the non-cohoused BCG-sc group, 3) the co-housed BCG-sc group and 4) the co-housed PBS-sc group. All mice were treated with PBS-sc or BCG-sc at PND 2, at which time the co-housing period began. At six weeks of age the co-housing period was terminated, and all groups were infected with *Mtb*-aerosol. After four weeks, pulmonary bacterial burdens and immunophenotyping were assessed. Both non-cohoused and co-housed BCG-sc groups had significantly reduced bacterial burdens post-infection compared to non-cohoused PBS controls (**Fig. 4f**). Interestingly, the co-housed PBS group displayed significantly reduced lung bacterial burdens compared to the non-cohoused PBS control (**Fig. 4f**), indicating that the co-housed BCG-microbiota afforded protection against infection. Immunophenotyping analysis revealed significantly increased levels of γδ T cells in the lung (**Fig. 4g**), predominantly the Vγ6^+^ subset (**Fig. 4h**, **Extended Data Fig. 5j-l**). This corroborated with an increased production of IL-22 but not IL-17A (**Extended Data Fig. 6m-n**). There were no changes in αβ T cells, including CD4^+^ and CD8^+^ subsets and NK cells (**Extended Data Fig. 5o-q**). The presence of BCG was not detectable by qPCR analysis within the fecal samples of any of the groups compared to purified culture positive controls (**Extended Data Fig. 5r**). Taken together, neonatally administered BCG-sc altered the intestinal microbiome in a unique manner, which was necessary for the expansion of lung γδ T cells and lowering pulmonary bacterial burdens, further supporting the role of the gut-lung axis in BCG-mediated γδ T cell protection against TB.

### Human Neonatal BCG Vaccination Expands **γδ** T Cells With Enhanced Antimicrobial programming

To complement the murine model of BCG vaccination, paired blood samples were obtained from a cohort of Singaporean infants (N=5) at birth and three months after receiving intradermal BCG (BCG-id) vaccination, which was administered at birth in accordance with the Singaporean national immunization schedule^26^ (**Fig. 5a**). Peripheral blood mononuclear cells (PBMCs) were isolated and subjected to single-cell RNA sequencing (scRNA-seq) and T cells were investigated. **Fig 5b** shows uniform manifold approximation and projection (UMAP) plot of total T cells, distinguishing CD4^+^, CD8^+^ and γδ T cells. Notably, the frequency of γδ T cells was significantly increased at 3-months (HB3) compared to birth (HB1), (paired t-test; p-value = 0.0448; **Fig. 5c**). Differential gene expression analysis comparing the transcriptome of γδ T cells at the two time points identified 143 differentially expressed genes (DEGs), of which 113 were upregulated and 30 were downregulated at 3 months of age (**Fig. 5d**, **Table S1**). Upregulated genes included IL32, GZMA, OAS2, KLRG1, PRF1, which encode cytolytic molecules important in host defense, indicating enhanced activity of γδ T cells at 3-months of age.^19, 27, 28^ Gene ontology (GO) analysis for biological processes further revealed enrichment of pathways related to T cell-mediated cytotoxicity and antigen processing and presentation among the upregulated genes at 3 months of age (**Fig. 5e**). Based on the expression of TRDV1 and TRGV9 genes, human γδ T cells were further divided into Vδ1 and Vδ2 subsets (**Fig 5b**, **Extended Data Fig. 6a**). Among these, Vδ2 cells showed a trending increased frequency at HB3 compared to HB1 (**Extended Data Fig. 6b**). DEG analysis between HB3 and HB1 showed several common genes among Vδ1 and Vδ2 subsets such as KLF6, PFN1, CYBA, JUNB, FTH1 (**Extended Data Fig. 6c,d**). GO analysis revealed enrichment of pathways related to response to virus and T cell activation among DEGs of Vδ1 and Vδ2, respectively (**Extended Data Fig. 6e**) Together, these findings suggest that, as in mice, BCG-id vaccination in human infants promotes the expansion of γδ T cells. In addition, BCG-educated γδ T cells in human infants are transcriptionally altered with a gene expression profile compatible with an increased capacity to induce protection.

## Discussion

The mechanism of protection of the BCG vaccine has puzzled the scientific community for over a century, as neonatal but not adult BCG-id vaccination offers substantial protection against TB disease^3^. Herein, we have demonstrated that dissemination of BCG due to the unique skin structure of neonates leads to the expansion of γδ T cells via the gut-lung axis, to provide protection against *Mtb* infection.

Early studies from the 1970s have demonstrated that systemic BCG vaccination (BCG-iv) in adult NHPs provided significantly greater protection against *Mtb* infection than BCG-id vaccination^29, 30^. More recently, BCG-iv vaccination in NHPs was shown to prevent *Mtb* infection altogether and induce sterilizing immunity^4, 31^. In an adult mouse model of TB, we also reported that BCG-iv vaccination reprograms HSCs in the bone marrow, biasing myelopoiesis and promoting the generation of memory monocytes/macrophages with enhanced anti-*Mtb* activity^6^. Here, we demonstrate that cutaneous BCG vaccination in neonatal mice can mimic the systemic protective effects of BCG-iv vaccination in adults. However, the mechanism of protection differs: in neonates, protection is mediated primarily by γδ T cells rather than monocytes/macrophages. Unlike BCG-iv vaccination in adults, neonatal BCG vaccination biases HSCs toward lymphopoiesis rather than myelopoiesis (**Fig. 2c-d**). Reprogramming of HSCs may underlie the durable immune protection conferred by early life vaccination against *Mtb*. Although we have not yet examined the transcriptomic and epigenetic reprogramming of hematopoietic stem and progenitor cells in neonates, it is tempting to speculate that BCG imprints a unique program in CLPs before their thymic trafficking, thereby enhancing γδ T cell development. This hypothesis requires further investigation.

Consistent with our findings, following BCG vaccination, γδ T cells expand and serve as major IFN-γ producers in infants and children^21^. They also play key roles during both acute^15^ and chronic^19^ responses to *M. tuberculosis* infection by producing IL-17^15, 32^. In patients with active TB, γδ T cells display an increased capacity for IL-17 production^15, 18, 32^. Furthermore, several studies using diverse models have documented the expansion and cytokine responses of γδ T cells following mycobacterial infection^15, 32, 33, 34, 35, 36^, with emerging mechanistic links to the gut microbiome^20, 37, 38^. The principle of the BCG-associated microbiome signature has been demonstrated in several key studies^37, 38^. In one study using an adult BCG vaccinated cohort, the variability in the measured cytokine response was associated with distinct microbial communities and metabolite production, for example the *Roseburia* genus and phenylalanine metabolism^37^. The interface between the microbiome and immune functions of mucosal-associated invariant T (MAIT) cells have recently garnered much interest. In one study, MAIT cell abundance and function, including granzyme B production, was linked with specific gut microbes in a cohort of active TB household contacts^20^. Further, MAIT cell function has been linked to early-life signals; colonization of key intestinal commensals later in life could not replicate the imprinting of MAIT cells in early-life, mediated via MAIT cell recognition of microbial-derived riboflavin^25^. In the present study, BCG-sc vaccination enriched *Prevotellaceae*, *Tannerellaceae*, and *Bifidobacteriaceae* microbial families compared to controls. Interestingly, the genomes of *Prevotellaceae* and *Tannerellaceae* taxa typically encode all essential functional genes involved in riboflavin biosynthesis^39^, and in another study, both taxa encoded the most abundant riboflavin synthesis genes^25^.

Further, in a recent work, responses to early-life antibiotics in infants linked the presence of *Bifidobacteriaceae* with improved responses and higher vaccine antibody titres^13^.

The indirect expansion of neonatal γδ T cells mediated by the BCG microbiome signature is conceivable given several key observations presented here and by other groups: 1) BCG was not detectable within the fecal DNA slurry, implicating an indirect mechanism of action, 2) γδ T cell antigen recognition is MHC-independent and includes microbial metabolites^40^, 3) γδ T cells are highly abundant during early-life within the lungs and gut^10, 11, 41^, which are colonized by the microbiota, and 4) the early dialogue between microbiota and immune cells is essential for appropriate unconventional T cell programming and function^25^. Further studies are required to carefully map the kinetics and functions of γδ T cells post-vaccination and through host development, as well as their capacity for diverse microbial metabolite recognition. Thus, early-life γδ T cells and microbiome present as pivotal and early indicators of effective BCG-mediated protection against *Mtb* infection, which should be leveraged in future vaccine and therapeutic strategies. This investigation, among an emerging body of works, demonstrates the importance of systemic BCG vaccine dissemination in the dialogue and development of the innate and adaptive arms of immunity against pulmonary pathogens.

A major barrier to the development of effective TB vaccines is the limited understanding of the mechanisms that confer protection against *Mtb* infection or prevent progression to active disease. The BCG vaccine remains the safest and most widely used TB vaccine, having been administered to over 100 million infants within the first week of life. Importantly, neonatal BCG vaccination can systemically disseminate^42^, conferring protection against pulmonary TB and extrapulmonary disease. Beyond TB, BCG is the only approved immunotherapy for non-muscle invasive bladder cancer^2^. We and others have recently demonstrated that its efficacy depends on systemic dissemination, especially to the BM, where it reprograms HSCs in both preclinical and clinical settings^6, 7, 43^. Notably, in the 1970s, intravenous administration of BCG (2×10 –3×10 CFU) was tested in 19 patients with advanced melanoma who were unresponsive to chemotherapy, producing remarkable inhibition of tumor growth^44^. In these immunocompromised patients, toxicity was tolerable at doses below 10 CFU^44^. Moreover, systemic BCG dissemination following neonatal vaccination has been associated with only rare symptomatic cases (e.g., fever, jaundice)^42^, and BCGosis remains uncommon^45^ even in immunocompromised patients with bladder or skin cancer. Taken together, these findings suggest that systemic BCG administration is a critical determinant of protection and prevention of *Mtb* infection, as shown in non-human primates^4^. Furthermore, virtually all current TB vaccine candidates in clinical trials target conventional T cells and focus on inducing type II IFN responses. However, the success of this strategy has been limited. Here, we demonstrate that unconventional T cells—with their unique functional capacities, including the production of IL-17 and IL-22, may represent a far more potent target for developing a novel TB vaccine. Given that *M. tuberculosis* continues to cause 1.5 million deaths annually, it is both feasible and timely to explore alternative vaccination strategies.

## Methods

### Mice

The following mice were purchased from Jackson Laboratories: C57BL/6, γδTCR^-/-^ (B6.129P2-Tcrdtm1Mom/J), and IL-17^-/-^ (B6.129(SJL)-Il17atm1.1(icre)Stck/J). IL-22^-/-^ mice were generously provided by Genentech. Mice used for neonatal studies were treated at post-natal day (PND) 2 or 3. Adult mice were treated at 6-8 weeks of age. Experiments were performed using both female and male age- and sex-matched mice. All animals were housed and inbred at the animal facility of the Research Institute of McGill University under specific pathogen-free conditions with ad libitum access to food and water, a temperature of 21°C (±1 °C), relative humidity of 40-60% (± 5%) and light cycle of 12h on, 12h off (daily cycle). All experiments involving animals were approved by the McGill University Animal Care Committee (permit no. 2010-5860) in accordance with the guidelines set out by the Canadian Council on Animal Care.

### Bacterial culture

*Mtb* H37Rv and BCG-TICE were grown in 7H9 broth (BD) supplemented with 0.2% glycerol (Wisent), 0.05% Tween80 (Fisher), and 10% albumin-dextrose-catalase (ADC) under constant shaking at 37°C. For *in vivo* experiments, *Mtb* (H37Rv) bacteria or BCG-TICE in log growing phase (OD 0.4 – 0.9) were centrifuged (4000 RPM, 15 minutes) and resuspended in sterile PBS. Single cell suspensions were obtained by passing the bacteria 10-15 times through a 22G needle (Terumo).

### *Mtb* infection or BCG vaccination of mice

Throughout the study, adult mice were subcutaneously (sc) or intravenously (iv) vaccinated with BCG-TICE with a dose of 1x10^6^ single-suspended bacteria in 100uL PBS, unless otherwise indicated. Neonatal mice were were subcutaneously (sc) or intrahepatically vaccinated with BCG-TICE with a dose of 1x10^6^ single-suspended bacteria in 50uL PBS. For aerosol infection, mice were infected with approximately 50-100 CFU of *Mtb* H37Rv in a nose-only aerosol exposure unit (Intox). Infection dose was confirmed by enumerating the lung CFU one-day after infection.

### Histopathological analysis

Lungs were inflated and fixed for 48 h with 10% formalin, and then embedded in paraffin. Sections (5 μm) were cut and stained with haematoxylin and eosin. Slides were scanned at a resolu-tion of x20 or ×40 magnification, and pictures were taken using a Leica Aperio slide scanner (Leica). Quantification of hematoxylin and eosin (H&E)-stained slides was performed using ImageJ software (National Institutes of Health (NIH), as in Tran *et al*. 2024^5^.

### Mycobacterial CFU Enumeration

For CFU enumeration in tissues, organs were homogenized in 1ml 7H9 broth (BD) supplemented with 0.2% glycerol (Wisent), 0.05% Tween80 (Fisher), and 10% ADC using OmniTip Plastic Homogenizer Probes (Omni International). Serial dilutions in PBS+0.05% Tween80 were plated on 7H10 agar plates with 10% OADC enrichment and PANTA (BD). For differentiation of *Mtb* CFU of BCG-vaccinated mice, samples were plated on *Mtb-*selective 7H10 agar plates with 3ug/L of thiophene-2-carboxylic acid hydrazide. Plates were then incubated at 37°C and counted after 21 days.

### Cohousing Study

Sex-matched mice at PND 2-3 were divided into 3 groups: 1) PBS-sc, 2) BCG-sc or 3) Co-housed. All animals were housed in the same room, and originated from the same breeding source. All neonates were treated with PBS-sc or BCG-sc in groups 1 and 2, respectively, and were weaned after PND 21. In group 3, at PND 2-3, in each cage 2-3 neonates received PBS-sc while 2-3 neonates received BCG-sc (as indicated by a front paw tattoo), keeping a maximum of 5 mice per cage. Mice were weaned at PND 21, but co-housing was maintained until PND 42 (6 weeks of age). At 6 weeks of age, all mice were infected with *Mtb*-aerosol as described above. Following 6 weeks of infection, all mice were euthanized and organs were collected for CFU enumeration and flow cytometry analysis. Fecal samples were collected from mothers prior to neonate birth, and from neonates prior to and post-*Mtb* aerosol infection.

### Flow Cytometry

Lung tissues were collected and minced before collagenase IV digestion (100 U) for 1 h at 37 °C. Lungs were filtered through a 70 μm nylon mesh, and red blood cells were lysed. Bone marrow was obtained by cutting the ends of tibiae and femurs and spinning down both bones for 5 seconds in a microtube. Samples were spun down and red blood cells were lysed. For cytokine measurement in lung γδ T cells, cells were incubated for three hours at 37 °C under stimulated or unstimulated conditions in RPMI media supplemented with FBS (10% v/v), Pen/Strep (Gibco, 1% v/v), HEPES (0.015 M), β-ME (Gibco, 1:750) and containing GolgiStop (BD, 1:1500). Under stimulated conditions, cells were treated with media also containing IL-1β (0.1 μL/mL) and IL-23 (0.2 μL /mL). After incubation, the cells were washed with PBS prior to staining.

Cells were stained with viability dye eFluor-506 (Invitrogen; 1:1000 dilution) for 20 minutes at 4°C. Subsequently, the cells were washed with PBS supplemented with 0.5% BSA (Wisent) and incubated with anti-CD16/32 (BD; 1:200) at a concentration of 1:200 in PBS/0.5% BSA at 4°C for 10 minutes, except for LKS, HSCs, MPPs and downstream progenitors staining. Cells were then stained for extracellular markers for 30 min at 4°C. Identification of adaptive immune cells was performed using BUV737-conjugated anti-CD45.2 (1:200), FITC-conjugated anti-CD3 (1:200), APC-Cy7-conjugated βTCR (1:200), PerCP-conjugated anti-γδTCR (1:200), eFluor450-conjugated anti-CD4 (1:200), BV650-conjugated or APC-conjugated anti-CD8 (1:200) and BUV395-conjugated anti-NK1.1 (1:200). For TCR-Vγ chain characterization of γδ T cells, cells were additionally extracellularly stained with PECy7-conjugated anti-Vγ2, PE-conjugated anti-Vγ3, and APC-conjugated anti-Vγ1. For Vγ6 staining, the monoclonal antibody specific for TCR-Vγ6 was kindly provided by Dr. Robert Tigelaar (Yale School of Medicine, Yale University). The culture supernatant of the anti-Vγ6 IgM secreting-17D1 hybridoma cells was directly added to each well for staining (50 μL for 30 minutes at 4 °C), followed by incubation with a secondary BV421-conjugated rat anti-mouse IgM antibody (1:150, 30 minutes at 4 °C). For intracellular staining, cells were fixed and permeabilized using the FOXP3 Transcription Factor Staining Kit (eBioscience) for 1 hour at 4 °C. Cells were further stained for 1 hour at 4 °C, for the assessment of cytokine production. Cytokine staining included: PE-conjugated anti-IL-22–PE (1:100), BV605-conjugated anti-IL-17A (1:100), BV786-conjugated RORγt (1:100), BV421-conjugated anti-IL-13 (1:100), BV786-conjugated anti-IL-4 (1:100), and APC-conjugated anti-IFNγ (1:100). Blood was obtained through cardiac puncture and directly stained with the same conjugated antibodies as listed above, after which cells were lysed. For staining of BM LKS, HSCs, MPPs and progenitors: Streptavidin–APC-Cy7 (1:100), anti-c-Kit–APC (1:100), anti-Sca-1–PE-Cy7 (1:100), anti-CD150– eFluor450 (1:100), anti-CD48–PerCP-eFluor710 (1:100), anti-Flt3–PE (1:100), anti-CD34–FITC (1:100), anti-CD16/32 PerCP-eFluor710 (1:100) and anti-CD127 BV786 (1:100) were added and incubated at 4°C for 30 minutes. Flow cytometry was performed using a BD LSR Fortessa X-20 (BD Biosciences) with FACSDiva version 8.0.1 (BD Biosciences). Analysis was performed using FlowJo version 10.10.0.

### Single-cell RNA-seq data generation

See Chen et al, for the cohort details and sample collection and processing for storage (reference). The longitudinal single-cell RNA sequencing (scRNA-seq) data from PBMCs of infants (n=5) at birth and 3 months of age was utilized for mining γδ T cells.

### Single-cell RNA-seq data analyses

Raw FASTQ files were aligned to the human reference genome (GRCh38) using Cell Ranger (v7.1.0). The resulting data were processed with the Seurat R package (v4.4.0). After removal of low-quality cells, doublets were identified and excluded using DoubletFinder (v2.0.3). Dimensionality reduction was performed via PCA followed by UMAP, and unsupervised clustering was conducted using Seurat’s FindClusters function. The γδ T cells were identified as CD3^+^ cells expressing TRDV1 or TRGV9. Their frequencies were calculated in each sample, and changes between birth and 3 months were assessed using a paired t-test after confirming normality with Shapiro-Wilk test. DEGs were identified using Seurat’s FindMarkers function with parameters logfc.threshold = 0 and min.pct = 0. Genes with adjusted p-values < 0.05 were considered significant, and volcano plot was generated using EnhancedVolcano package (v 1.24.0). GO enrichment analysis was performed using clusterProfiler package (v4.2.0), with all expressed genes (n=21,678) from the integrated Seurat object used as background. Gene identifiers were converted to ENTREZ IDs using the bitr function and the org.Hs.eg.db annotation package (v3.14.0). GO enrichment analysis was conducted separately for up- and down-regulated DEGs at 3 months vs. birth using compareCluster function with fun = "enrichGO", ont = "BP", pvalueCutoff = 0.05, and qvalueCutoff = 0.10.

### Bacterial DNA extraction and 16S rRNA gene sequencing

Mouse fecal pellets were collected and stored at -80C. Bacterial DNA was extracted from fecal pellets using the QIAGEN DNeasy PowerSoil Pro Kit per the manufacturer’s instruction. The V4 hypervariable region of the 16S rRNA gene was amplified with primers Forward Primer (515F): (GTGCCAGCMGCCGCGGTAA) and Reverse Primer (806R): (GGACTACHVGGGTWTCTAAT) . Amplicons were sequenced using an Illumina MiSeq sequencer, generating 2 x 250 length bp paired-end reads in two independent experiments and sequencing runs.

Primers and adapters were removed from sequences prior to analyses. Qiime2 (v. 2025.7)^46^ was used for quality control and processing. DADA2^47^ was used to trim low quality bases, join forward and reverse pairs, and remove chimeric reads. Taxonomy was assigned to amplicon sequence variants (ASVs), using the Qiime2 feature classifier^48^. A full-length 16S rRNA gene pre-trained Naïve Bayes classifier (Silva v. 138) was used to assign taxonomy. Downstream analyses were then performed in R using Phyloseq (v. 1.50)^49^.

To perform beta-diversity analyses, libraries were first normalized by rarefying at 4,229 counts per sample using Phyloseq (v. 1.50)^49, 50^. Bray-Curtis distances were calculated using vegan (v.2.7-1)^51^. PERMANOVA (pairwiseadonis, v. 0.4.1) was used to assess significant differences between treatment groups in each independent experiment. P-values were adjusted using the Benjamini-Hochberg method. Beta-dispersion was performed (vegan, v.2.7-1) to determine that there were no differences in treatment variance. DESeq2 (v. 1.46) was used to determine differentially abundant bacterial families between treatment groups^52^. The experiment from which each sample originated was included as a covariate in the formula: ∼ experiment + treatment. Log_2_ fold-change > 1 or < -1 and p ≤ 0.05 were used as thresholds for significance. Enhanced-volcano (v.1.24) was used to visualize differentially abundant families.

### Bacterial RT-qPCR

cDNA was generated by qPCR using BrightGreen SYBR Green (ABM). Cq values obtained on the CFX96 PCR System (Bio-Rad) were analyzed using the 2^−ΔCq^ formula normalizing target gene expression to *SigA*. Primer sequences for *SigA* were: 5’-TGCAGTCGGTGCTGGACA -3’ and 5’-CGCGCAGGACCTGTGAGC -3’. Primer sequences for *IS1081* were: 5’- GAT-CATGGCCAAAGAGCTCG -3’ and 5’-GAGTGCACACCTTGATCGC -3’.

### Confocal Microscopy

Skin samples were fixed overnight at 4°C in 1% paraformaldehyde and stained for CD31 to visualize endothelial cells. Confocal microscopy was performed with a LSM 880 confocal microscope equipped with a 20 × /0.8 NA Plan-Apochromat objective (Carl Zeiss Microimaging). Z-stacks of 32 µm were acquired and maximum intensity projections are shown. Scale bars, 25 µm.

### Whole-Mount Immunofluorescence

#### BCG-Wasabi Injection and Tissue Preparation

PND 3 pups and adult C57BL/6 mice (6–8 weeks old) were intradermally injected with 7x10^6^ CFU of BCG-Wasabi. The injection site was circled with a permanent marker to ensure accurate localization during tissue collection. Mice were euthanized 5 minutes post-injection, and the injection site was excised. Skin samples were fixed in 1% paraformaldehyde (PFA; Electron Microscopy Sciences) for 24h at 4 °C. Following fixation, tissues were washed in PBS and incubated in blocking buffer (5% normal goat serum, 0.3% Triton X-100 in 1× PBS) for 2 h at room temperature on a shaker. Samples were then incubated with anti-CD31-AF647 (BioLegend, cat. no. 102516) diluted in blocking buffer for 1 h at room temperature on a shaker, followed by overnight incubation at 4 °C. After staining, tissues were washed twice with PBS prior to imaging.

#### Microscopy

Imaging was performed using a Nikon CSU-X1 spinning disk confocal microscope. For vasculature analysis, five randomly selected fields of view (FOV) per biological replicate were acquired using a 20× water-immersion objective. For intravascular BCG quantification, five FOV were acquired along the injection site using a 40× water-immersion objective.

#### Image Analysis

Vasculature diameter and area were quantified using ImageJ (NIH). Vessel diameter was calculated as the mean of 40 evenly distributed measurements across each FOV. Vasculature area was calculated as the ratio of CD31 vascular area to the total FOV area. Intravascular BCG-Wasabi was quantified using Imaris (Oxford Instruments) software with the surface-to-spot distance function, applying a threshold of 0.5 µm. Three-dimensional reconstruction of intravascular BCG was performed in Imaris using representative images acquired with the 40× objective.

#### Ethics Statement

All experiments involving animals were approved by the McGill University Animal Care Committee (permit number 2010–5860) in accordance with the guidelines set out by the Canadian Council on Animal Care.

All experiments involving human research participants were approved by the National Health-care Group Institutional Review Board in Singapore under the title "Gestational Immunity for Transfer GIFT" (DSRB Reference Number: 2020/00483). Written informed consent was obtained from all participating mothers, and the study adhered to the ethical principles stated in the Declaration of Helsinki.

The study protocol was registered on clinicaltrials.gov (NCT04802278). Seven mothers who had recovered from antenatal COVID-19 were recruited. They were confirmed positive for COVID-19 via real-time reverse-transcriptase polymerase chain reaction (RT-PCR) on nasopharyngeal swabs during pregnancy. Recovery was defined as the resolution of clinical symptoms, along with two consecutive negative SARS-CoV-2 RT-PCR tests, 24 hours apart. Infants born to these mothers at 35-40 weeks gestation were included in the study. Comprehensive RT-PCR testing for COVID-19 on the placenta, nasopharyngeal, vaginal, amniotic fluid, umbilical cord, and breast milk at delivery returned negative results for all convalescent mothers. A control group of six mother-infant dyads was also enrolled, with control mothers having no COVID-19 symptoms and a negative SARS-CoV-2 IgG test at recruitment. Birth parameters were compared between SARS-CoV-2 exposed uninfected (SEU) and control infants using Fisher’s exact test for categorical variables and the Wilcoxon rank-sum test for continuous variables. No significant differences were found between the groups.

## Data Availability

Code used for data analyses can be found at: https://github.com/anshulsinha1/BCG-Neonate-Manuscript. Raw 16S rRNA gene sequencing files were deposited in the NCBI sequence read archive and can be found using the BioProject ID: PRJNA1358291.

## Statistics

Data are presented as the mean ± S.E.M. Statistical analyses were performed using GraphPad Prism v10 software (GraphPad). Unless stated, statistical differences were determined using one-way ANOVA followed by Tukey’s multiple-comparisons test or two-way ANOVA followed by Tukey’s multiple-comparisons test.

## Acknowledgements

The authors acknowledge technical help from Veronica Sandy from the RI-MUHC ARD; Fiona McIntosh from the RI-MUHC CL3 facility; Z.M. Cordova, G. Fontes; the staff at the McGill Centre for Microbiome Research; the staff at the Meakins Christie Laboratories; and the staff at the RI-MUHC Histopathology platform. M.D. is funded by Canadian Institute of Health Research (CIHR) Project Grant-203950, a Canada Research Chairs (CRC-2025-00096) in Trained Immunity, and a McGill Interdisciplinary Initiative in Infection and Immunity (MI4) Grant. M.D. holds the Strauss Chair in Respiratory Diseases and is a fellow member of the Royal Society of Canada. M.S. is supported by a Fonds de Recherche du Québec–Santé Award. The funders had no role in study design, data collection and analysis, decision to publish or preparation of the manuscript. Schematic models in were created using BioRender (https://biorender.com).

## Author Contributions

Conceptualization: M.S., N.K., and M.D. Methodology: M.S., N.K., Y.C., L.F.J., A.S., K.C., I.V., V.L., K.A.T., S.H.B., B.T., T.J.S., N.G., E.K., J.M.L., Z.A., C.M., J.K., J.X., M.K., R.X., M.G.N., I.K., L.B.B., A.T., A.S., M.D. Investigation: M.G.N., I.K., L.B.B., A.T., A.S., M.D. Funding acquisition: M.D. Project administration: M.S., N.K., M.D. Supervision: M.D. Writing original draft: M.S. and M.D. Writing, review and editing: M.S., N.K., Y.C., A.S., K.C., B.T., E.K., M.K., R.X., M.G.N., I.K., L.B.B., A.T., A.S., M.D.

## Competing Interests

The authors declare no competing interests.

**Correspondence and requests for materials** should be addressed to Maziar Divangahi.

## Notes

### Competing Interest Statement

The authors have declared no competing interest.

