## Extended Figures 1-6 for "Neonatal BCG Vaccination Engages the Vasculature to Elicit γδ T Cell–Mediated Protec-tion against Tuberculosis"

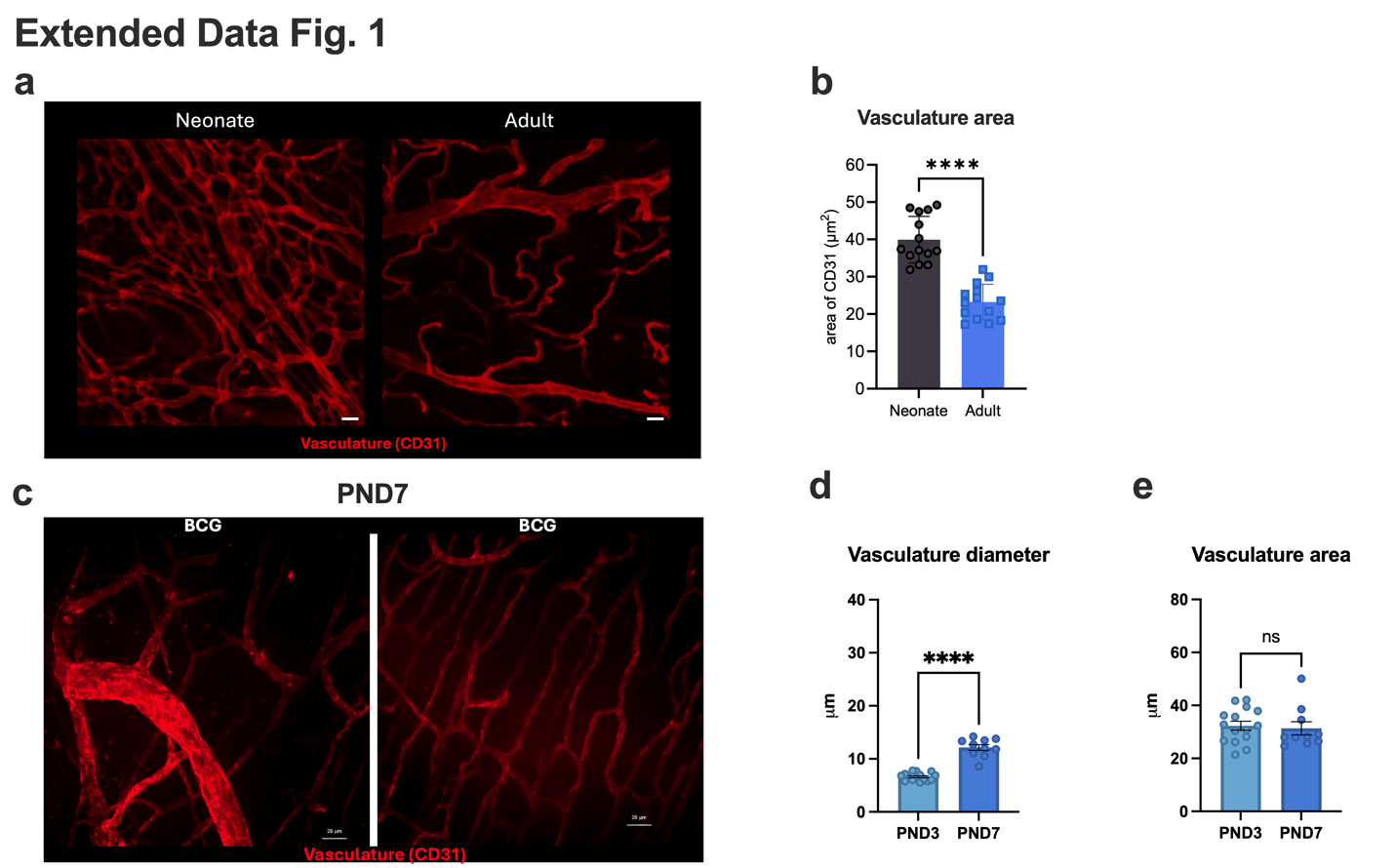

**Extended Figure 1: Three-dimensional reconstruction shows intravascular BCG only within neonatal skin vasculature. (a)** Representative confocal images with CD31 immunofluorescence labeling of dermal blood vessels in neonatal (PND3) and adult (6–8-week-old) mouse skin. **(b)** Quantification of blood vessels in neonatal and adult mouse skin quantified using ImageJ software from 3-6 fields of view from 3 mice per group. Statistical significance for comparison between neonates and adult skin was performed by an unpaired two-tailed t-test. Neonatal and adult mice were intradermally vaccinated with BCG-Wasabi and euthanized at various timepoints. Skin sections adjacent to the injection site were excised and stained for CD31 to visualize vasculature. **(c)** Two representative images from wholemount immunofluorescence staining of CD31^+^ vasculature in neonate skin with BCG-sc injections (at PND3). Mice were euthanized 4 days post-injection, at PND7 (Scale bar: 25 μm). **(d)** Average vessel diameter and (**e**) vasculature area was quantified at PND7, and compared with BCG-sc vaccination immediately post-injection at PND3.

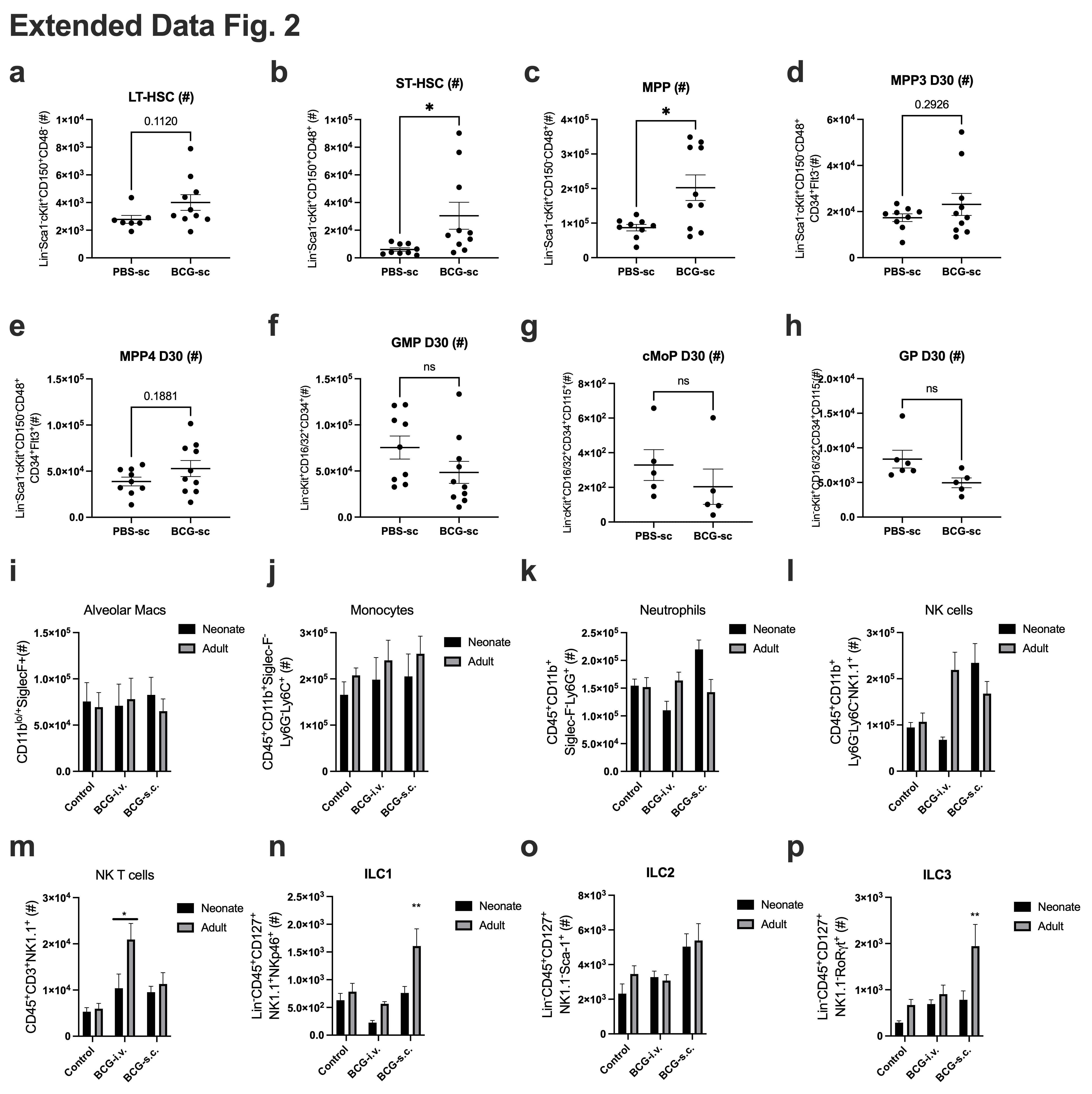

**Extended Figure 2: Neonatal BCG-sc induces LKS expansion, but not classical innate or adaptive cells.** Neonates were vaccinated with BCG-sc or PBS-sc control and BM was collected after 4 weeks. Absolute counts of **(a)** LT-HSCs, **(b)** ST-HSCs, **(c)** MPPs, **(d)** MPP3s, **(e)** MPP4s, **(f)** GMPs, **(g)** cMoPs, and **(h)** GPs. Neonates and adults were vaccinated with PBS-sc, BCG-iv and BCG-sc and lung tissue was collected after 8 weeks. The innate compartment was assessed by absolute counts of **(i)** alveolar macrophages, **(j)** monocytes and **(k)** neutrophils. Adaptive and innate lymphoid cells were also measured, including **(l)** NK cells, **(m)** NK T cells, **(n)** ILC1s, **(o)** ILC2s, and **(p)** ILC3s. Data are presented as mean ±SEM and are from one or pooled from two independent experiments. Data were analyzed using unpaired t-test or two-way ANOVA followed by Sidak’s multiple comparisons test.

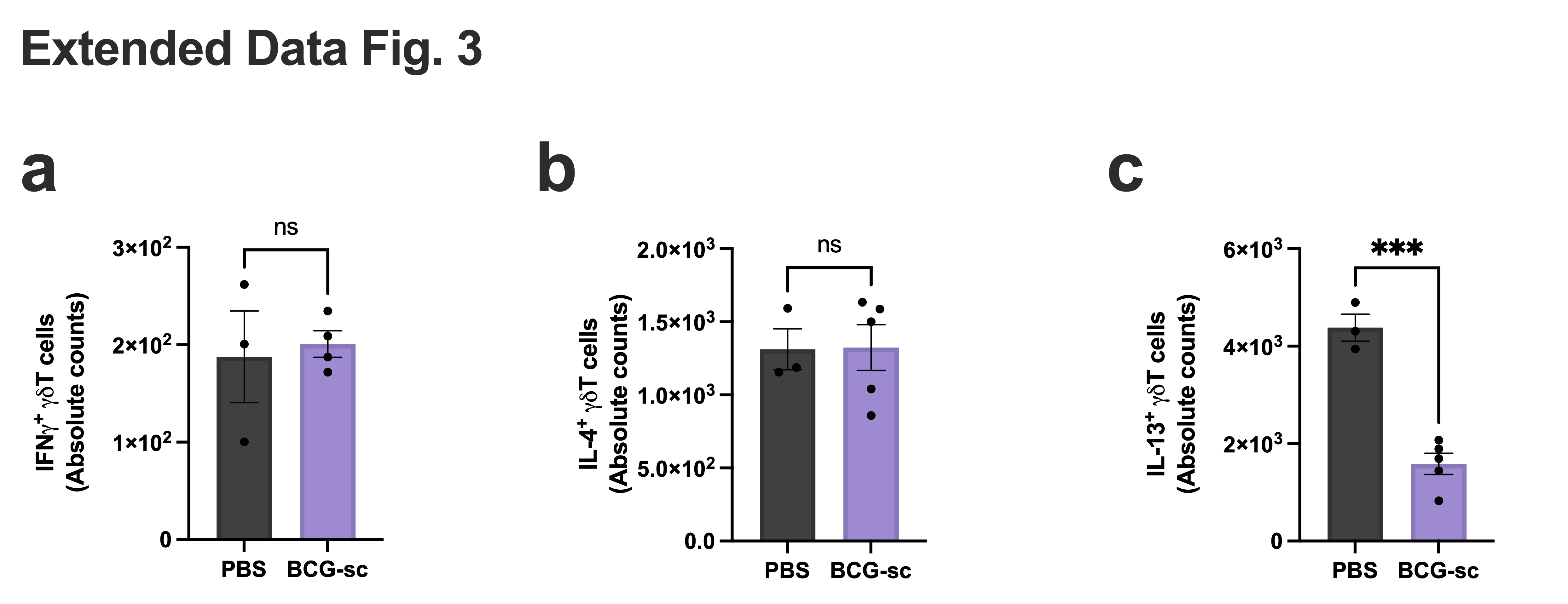

**Extended Figure 3: BCG vaccination does not induce type 1 and type 2 γδ T cells.** Absolute counts of **(a)** IFNγ^+^ γδ T cells, **(b)** IL-4^+^ 2 γδ T cells, and **(c)** IL-13^+^ γδ T cells after 4 weeks of PBS-sc or BCG-sc. N=3-5 per group. Data were analyzed using two-tailed unpaired t-test. Data are presented as mean ±SEM and are from one or pooled from two independent experiments.

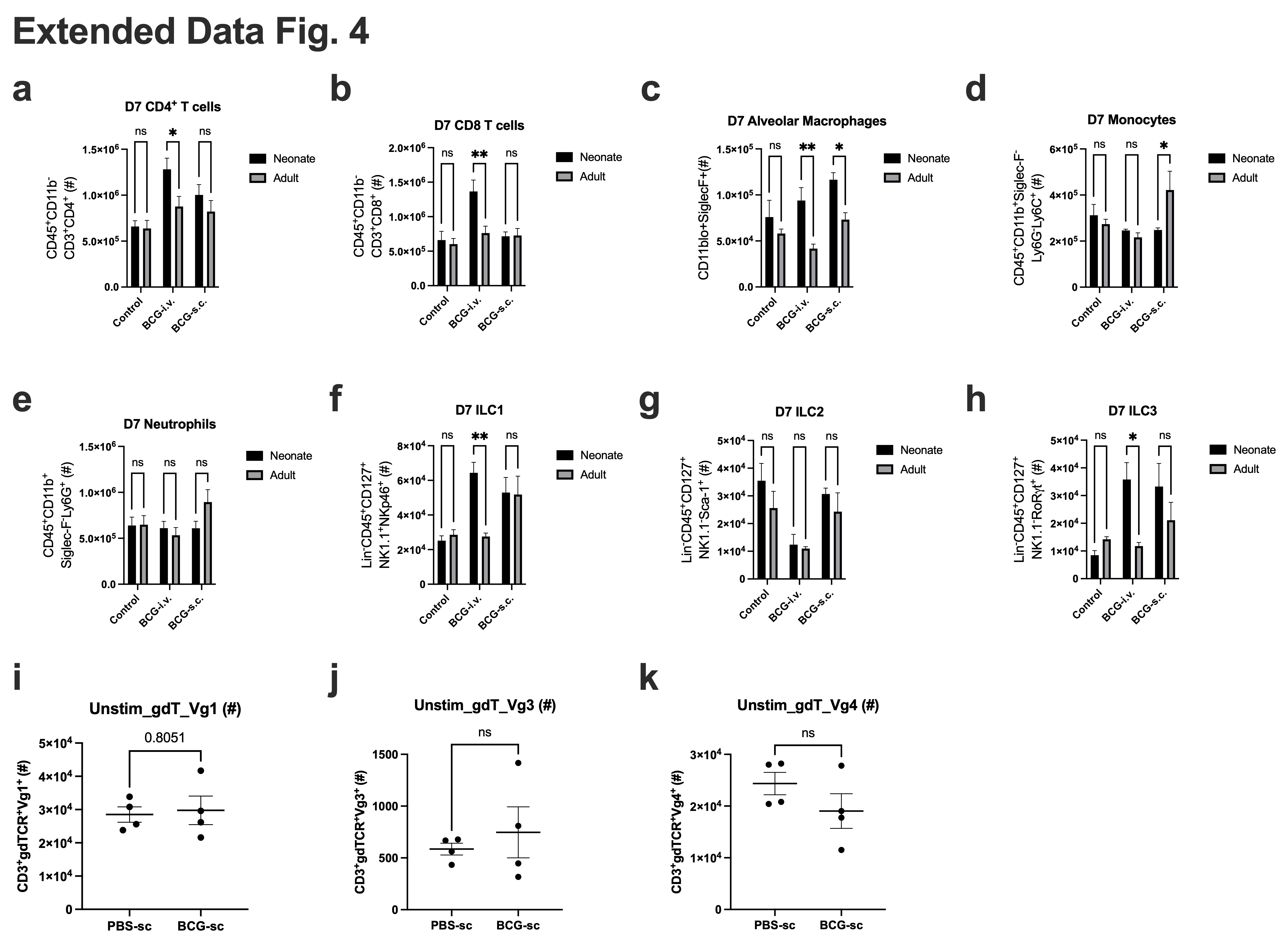

**Extended Figure 4: Lung immunophenotyping following *Mtb* infection and neonatal BCG vaccination.** Neonate or adult mice were vaccinated with PBS control, BCG-iv or BCG-sc and infected with Mtb-aerosol after 20 weeks. Lungs were collected for immunophenotyping after 7 days of infection. Absolute counts of **(a)** CD4^+^ T cells, **(b)** CD8+ T cells, **(c)** alveolar macrophages, **(d)** monocytes, **(e)** neutrophils, **(f)** ILC1s, **(g)** ILC2s and **(h)** ILC3s were assessed. Neonatal mice were infected after 6 weeks of vaccination, and lung tissue was collected after 4 weeks. **(i)** Tbet^+^ γδ T cells **(j)** Vγ4^+^ T cells **(k)** Vγ3^+^ T cells and **(l)** Vγ1^+^ T cells were measured (N=4 per group_. Data are presented as mean ±SEM and are from one or pooled from two independent experiments. Data were analyzed using unpaired t-test (**i-k**) or two-way ANOVA (**a-h**) followed by Sidak’s multiple comparisons test.

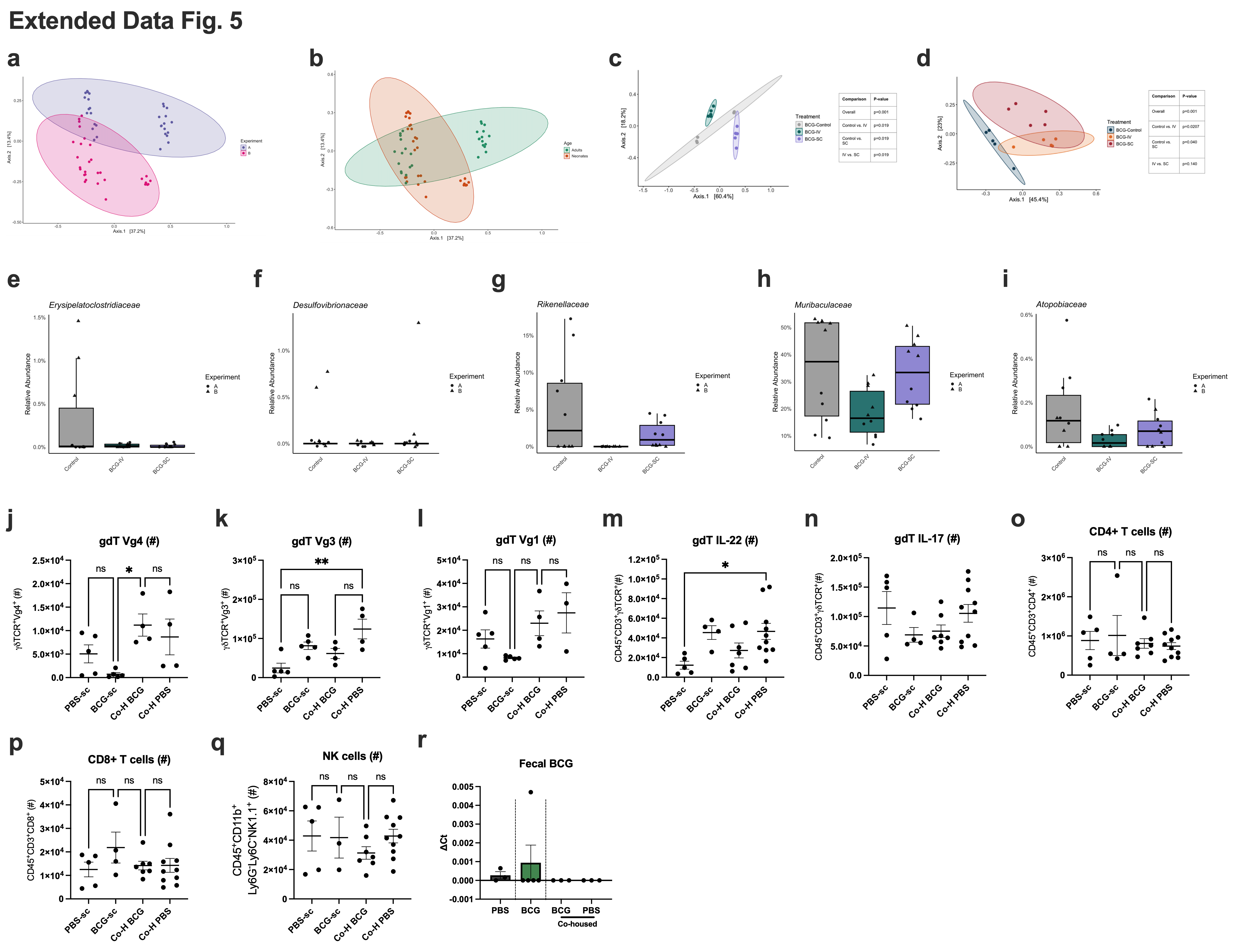

**Extended Figure 5: BCG co-housing induced a γδT17 immune profile**. Neonate and adult mice were administered PBS-sc, BCG-iv and BCG-sc and 8 weeks after vaccination fecal samples were collected for 16S rRNA gene sequencing analysis (N=5 per group). **(a)** Experiment- and **(b)** age-effects on bacterial diversity. PCoA were generated using Bray-Curtis dissimilarity. Principal Coordinates Analysis (PCoA) with Bray-Curtis dissimilarity for **(c)** neonatal mice or **(d)** adult mice. PERMANOVA was used to assess significance p < 0.05 between groups. These data are representative of the second of two experiments. Relative abundance of (**e**) *Erysipelatoclostridiaceae*, **(f)** *Desulfovibrioceae,* **(g)** *Rikenellaceae,* **(h)** *Muribaculaceae* and **(i)** *Atopobiacae* families in neonatal groups. Lung immunophenotyping was conducted post-BCG cohousing study. Absolute counts of **(j)** Vγ4^+^ T cells **(k)** Vγ3^+^ T cells and **(l)** Vγ1^+^ T cells were quantified (N=4-5 per group). (**m)** IL-22^+^ and **(n)** IL-17+ producing γδ T cells were also assessed (N= 5-10 per group). The adaptive compartment was measured including **(o)** CD4^+^ T cells, **(p)** CD8^+^ T cells and (**q)** NK cells (N= 5-10 per group). Data are representative of two independent experiments. **(r)** Fecal mRNA levels (ΔCt) of BCG insertion element IS1081^53^ relative to *SigA* housekeeping gene (N=3-5 per group). Data were analyzed using one-way ANOVA followed by Sidak’s multiple comparisons test and are presented as Mean ±SEM.

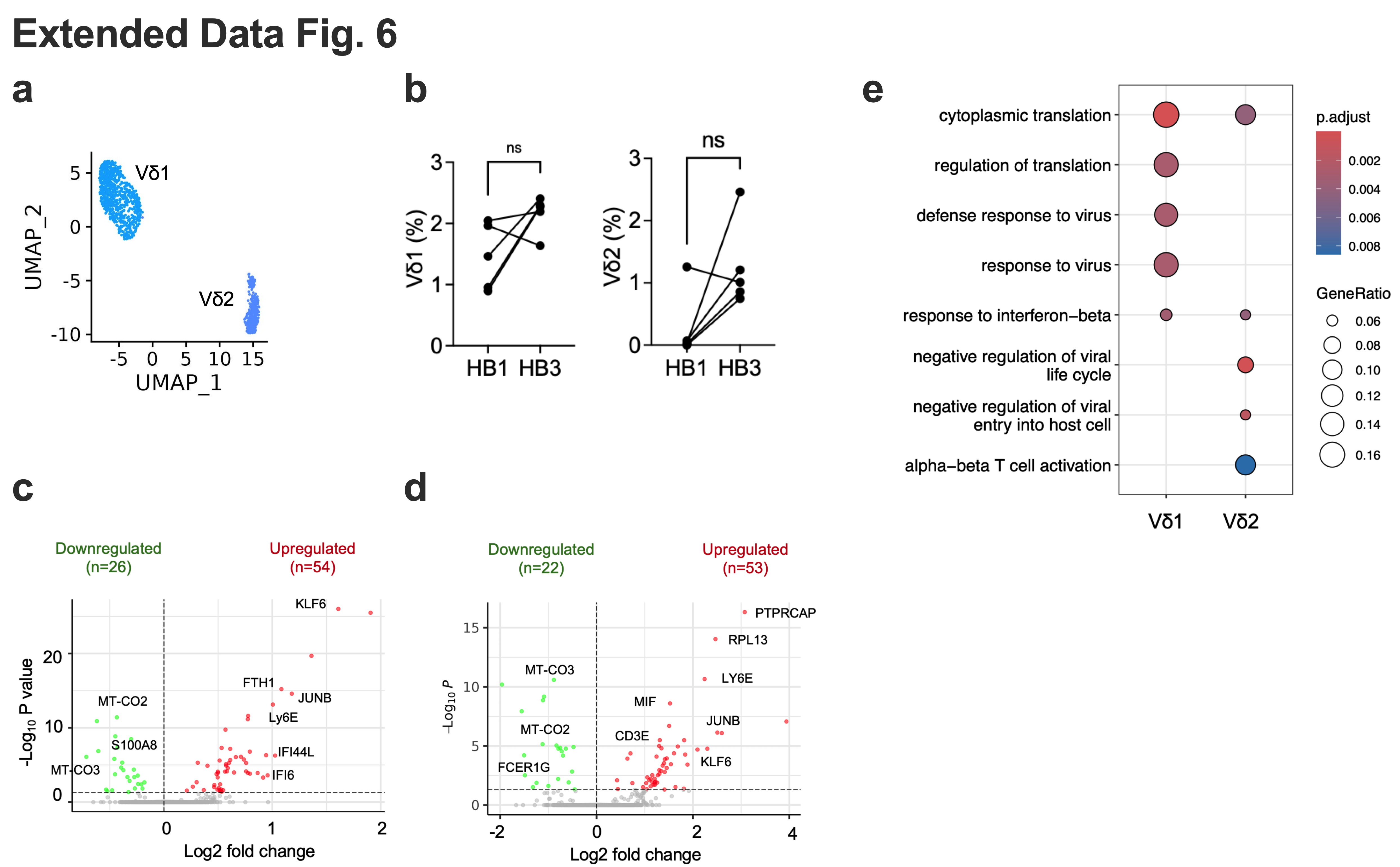

**Extended Figure 6:** **Transcriptional reprogramming in human neonatal γδ T cell subtypes.** **(a)** UMAP plot of Vδ1 and Vδ2 cells by single-cell RNA-seq in neonatal PBMCs. **(b)** Percentage of Vδ1 (left) and Vδ2 (right) cells in total PBMCs of infants at birth and at 3 months of age. Ns, not significant, paired t-test. **(c and d)** Volcano plot of DEGs in 3 months (HB3) vs birth (HB1) in Vδ1 (c) and Vδ2 (d) cells. Number of up- and down-regulated genes are indicated. **(e)** Enriched GO biological processes in 3 months (HB3) vs birth (HB1) DEGs identified in Vδ1 and Vδ2.

**Supplementary Information Legends**

**Supplementary Video 1: Neonatal skin consists of intricate networks of small vessels with many intravascular BCG colonies.**

Three-dimensional reconstructions of skin vasculature and fluorescent BCG generated using Imaris software for a representative neonatal mouse (at PND3) immediately following BCG vaccination.

**Supplementary Video 2: Adult skin sections display large vessels with rare intravascular BCG colonies.**

Three-dimensional reconstructions of skin vasculature and fluorescent BCG generated using Imaris software for a representative adult mouse immediately following BCG vaccination.

**Supplementary Video 3: Adult skin contains a vasculature-adjacent adipose tissue layer which entraps BCG.**

Representative video of adult skin vasculature, illustrating the spatial organization of tissue layers starting from adipose tissue to the vascular network.

**Supplementary Figure 1: Flow cytometry gating strategies for the evaluation of immune cells in the lungs of adult mice.**

**(a)** T cells and innate lymphoid cells (ILCs) were gated on single live CD45.2^+^ cells then further gated on CD3 positive or Lineage (CD5, CD8, CD4, CD19, CD45R, Ly6G/C, Ter119, CD11b) negative cells. CD3^+^ were further gated on NK1.1^+^γδTCR^-^ (NK T cells), NK1.1^-^γδTCR^+^ (γδ T cells), CD8^+^CD4^-^ (CD8^+^ cells), CD8^-^CD4^+^ (CD4^+^ cells). Lineage negative cells were further gated on CD127^+^ cells, with the expression of NK1.1^+^NKp46^+^ (ILC1), NK1.1^-^Sca1^+^ (ILC2), NK1.1^-^RORγt^+^ (ILC3). **(b)** Innate cells were gated on single live CD45.2^+^ cells then further gated on SiglecF and CD11b expression: SiglecF^+^CD11b^-^ (alveolar macrophages), SiglecF^+^CD11b^+^ (eosinophils). CD11b^+^ and SiglecF^-^ cells were further divided by Ly6G and Ly6C expression: Ly6G^+^Ly6C^+^ (neutrophils) and Ly6C^+^Ly6G^-^ (monocytes). Double negative (Ly6G^-^Ly6C^-^) cells were divided into dendritic cells (CD11c^+^F4/80^-^), macrophages (F4/80^+^NK1.1^-^) and NK cells (F/480^-^NK1.1^+^). **(c)** BM HSC staining was adapted from Khan *et al*.^7^, and was gated on single live cells and further gated on Lineage (CD5, CD4, CD8, CD19, CD45R, Ly6G/C, Ter119) negative cells. Lineage-committed progenitors were also gated on the expression of CD127, cKit and Sca-1: CD127^+^cKit^lo^Sca-1^lo^ (CLP), CD127^-^cKit^+^Sca-1^-^CD16/32^-^CD34^+^ (CMP), CD127^-^cKit^+^Sca-1^-^CD16/32^+^CD34^+^ (GMP), and CD127^-^cKit^+^Sca-1^-^CD16/32^-^CD34^-^ (MEP). Lineage negative cells were alternatively gated on cKit^+^ and Sca-1^+^ cells, and divided based on the expression of CD48 and CD150 into: MPPs (CD150^-^CD48^+^), ST-HSCs (CD150^+^CD48^+^), and LT-HSCs (CD150^+^CD48^-^). MPPs were further divided into MPP4 (Flt3^+^CD34^+^) and MPP3 (Flt3^-^CD34^+^) populations.

**Supplementary Table 1: List of differentially expressed genes in total, type 1 and type 2 γδ T cells from infants at 3 months of age.**
