## Supplementary figures and images for "Neonatal BCG Vaccination Engages the Vasculature to Elicit γδ T Cell–Mediated Protec-tion against Tuberculosis"

### Supplemental Figure 1

#### Supplementary Fig. 1

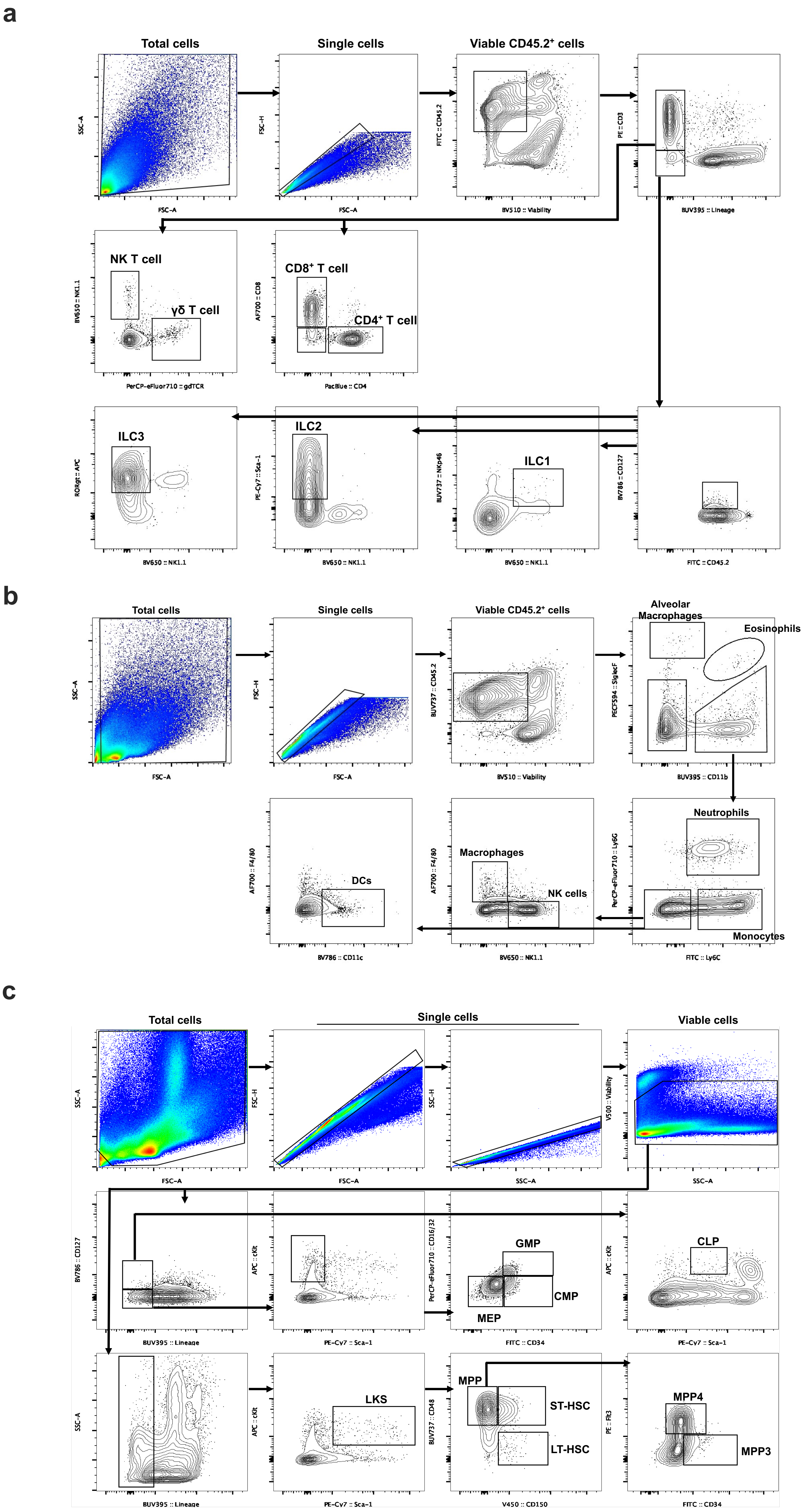
